# Chronic replication stress-mediated genomic instability disrupts placenta development in mice

**DOI:** 10.1101/2025.02.28.640689

**Authors:** Mumingjiang Munisha, Rui Huang, Jordan Khan, John C. Schimenti

## Abstract

Abnormal placentation drives many pregnancy-related pathologies and poor fetal outcomes, but the underlying molecular causes are understudied. Here, we show that persistent replication stress due to mutations in the MCM2-7 replicative helicase disrupts placentation and reduces embryo viability in mice. MCM-deficient embryos exhibited normal morphology but their placentae had a drastically diminished junctional zone (JZ). Whereas cell proliferation in the labyrinth zone (LZ) remained unaffected, JZ cell proliferation was reduced during development. MCM2-7 deficient trophoblast stem cells (TSCs) failed to maintain stemness, suggesting that replication stress affects the initial trophoblast progenitor pool in a manner that preferentially impacts the developing JZ. In contrast, pluripotency of mouse embryonic stem cells with MCM2-7 deficiency were not affected. Developing female mice deficient for FANCM, a protein involved in replication-associated DNA repair, also had placentae with a diminished JZ. These findings indicate that replication stress-induced genomic instability compromises embryo outcomes by impairing placentation.

## Introduction

Early embryonic development is characterized by rapid cellular proliferation and differentiation. In such a period, it is imperative for cells, especially stem cells that engender various components of the developing embryo, to maintain genomic integrity to promote proper formation, propagation and maintenance of cellular identity ^1,2^. Genome maintenance requires precise coordination of various cellular processes including those involved in DNA replication and repair ^3,4^. Disruptions to these processes can lead to genomic instability (GIN), a cellular state characterized by elevated DNA damage such as double-strand breaks (DSBs), cell cycle delay/arrest and mutation accumulation ^4^.

Most of our knowledge of basic genome maintenance mechanisms has been derived from studies on cancer cell lines. The impact of GIN upon organismal development has been focused primarily on the embryo proper. Although extraembryonic placental cells share several characteristics with cancer cells such as invasiveness and high levels of copy number variation ^5^, the mechanisms underlying genome maintenance in placental cells are relatively understudied. For example, mouse trophoblast giant cells (TGCs) are highly polyploid and selectively amplify regions containing key placental genes, causing extensive copy number variation throughout the genome ^6–8^. Differentiated trophoblasts are resistant to GIN as they do not activate DNA damage checkpoints in response to genotoxic stress ^9–12^. This is understandable given the transient nature of the placenta. Similarly, studies using term human placental tissues have revealed that high mutation rates, genome amplification and senescence are common features of human trophoblast cells ^13–15^. Mutations in genes involved in genome maintenance can lead to diseases ^3^. Though such mutations are rare, mammalian embryos and placentae face unique challenges during normal development that can impose GIN ^1^. While excessive GIN can result in adverse embryo outcomes by negatively affecting placental function via inducing senescence and inflammation ^16,17^, the extent to which GIN is tolerated at different stages of placenta development needs further investigation.

Using a unique mouse model with high intrinsic GIN, known as *Chaos3 (C3)*, we showed that deficiencies in the DNA replicative helicase can result in sex- and parent-of-origin dependent embryonic semi-lethality ^17^. The *C3* allele is a hypomorphic missense point mutation in the minichromosome maintenance complex 4 (*Mcm4)* gene which encodes a protein that is part of the core catalytic unit of the CMG (CDC45, MCM2-7, and GINS complex) replicative helicase ^18^. MCMs are loaded as heterohexamers of MCM2-7 onto DNA replication origins (sites of replication initiation) during the G1 phase of the cell cycle, a process referred to as origin “licensing” ^19,20^. Only a portion of licensed origins initiate bidirectional replication during S-phase, leaving the rest to serve as dormant origins (DO). These DOs are crucial when cells experience replication stress (RS), enabling rescue of chromosomal regions left unreplicated from stalled or collapsed replication forks ^19,20^. The amino acid change in MCM4^C3^ disrupts its interaction with MCM6, destabilizing the MCM2-7 complex. This leads to persistent RS and activation of p53-dependent upregulation of miRNAs that target *Mcm* mRNAs, reducing DOs, increasing hypersensitivity to DNA damaging agents, and dramatically increasing cancer susceptibility ^18,21,22^.

*Mcm4^C3/C3^* mice are viable and fertile, and exhibit elevated micronuclei (a nuclear membrane-bound cytosolic DNA) in erythrocytes, a hallmark of persistent RS and GIN ^18^. Genetic depletion of another MCM in the *Mcm4^C3/C3^* mice (for example with the genotype *Mcm4^C3/C3^ Mcm2^Gt/+^*; “*Gt*” is a gene trap null allele) resulted in embryonic semi-lethality in which 40% of males survive to birth compared to only 15% of females ^17,22^. The female lethal bias was associated with elevated markers of placental inflammation, and the sex skewing could be eliminated by suppressing inflammation pharmacologically ^17^. We hypothesized that the inflammation and female-biased embryonic lethality was caused by activation of innate immune pathways that sense cytosolic nucleic acids. Here, we report that key cytosolic DNA sensors (cGAS-STING; RIG1-MAVs) are not responsible. Rather, elevated RS causes defective trophoblast stem cell (TSC) proliferation/maintenance in mutants with high levels of GIN resulting in defective placentation, and inflammation is likely a consequence of this disruption. Overall, these studies underscore the sensitivity of placentation to RS-induced GIN during early development.

## Results

### Persistent replication stress (RS) leads to a small placenta and intrauterine growth restriction

As mentioned earlier, deficiencies in DNA replicative helicase causes embryonic semi-lethality, preferentially affecting female embryos ^17^. Female-biased embryonic lethality was observed in mice bearing the semi-lethal genotype *Mcm4^C3/C3^ Mcm2^Gt/+^* from the sex-skewing mating pairs, where the dam genotype is *Mcm4^C3/C3^* and the sire genotype is *Mcm4^C3/+^ Mcm2^Gt/+^* (Fig. 1A). However, this female bias was abolished in the reciprocal mating where the dam and sire genotypes were switched (Fig. 1A). We previously proposed that placental inflammation might be the culprit of female-biased embryonic semi-lethality for the following reasons: 1) RNA-Seq of E13.5 placenta revealed an enrichment of inflammatory signatures; 2) anti-inflammatory (Ibuprofen or testosterone) treatment of pregnant dams rescued the female-bias; and 3) deficiency for the anti-inflammatory gene *Il10rb* was synthetically lethal with *Mcm4^C3/C3^*, but viability could be rescued by ibuprofen treatment of pregnant dams ^16^. Given that elevated micronuclei is the hallmark of *Mcm4^C3/C3^* mice ^18^, we hypothesized that a cytosolic nucleotide sensing pathway, such as cGAS-STING, was responsible for increased placental inflammation, and that male embryos were preferentially protected by high levels of testosterone around the time of death. To test this hypothesis, we genetically deleted *Tmem173* (encoding STING protein in the cGAS-STING pathway) and separately *Ddx58* (encoding RIG1 protein in the RIG1-MAVS pathway) in the *Mcm4^C3/C3^ Mcm2^Gt/+^* background. Deletion of neither of these genes rescued female-biased embryonic lethality (Supplementary Table 1). Additionally, knocking out *Myd88*, encoding an obligate co-factor for the Toll-Like-Receptor signaling, in the *Mcm4^C3/C3^* background was synthetically lethal post-wean (not shown), not permitting us to generate animals of the desired genotype.

**Figure 1.**
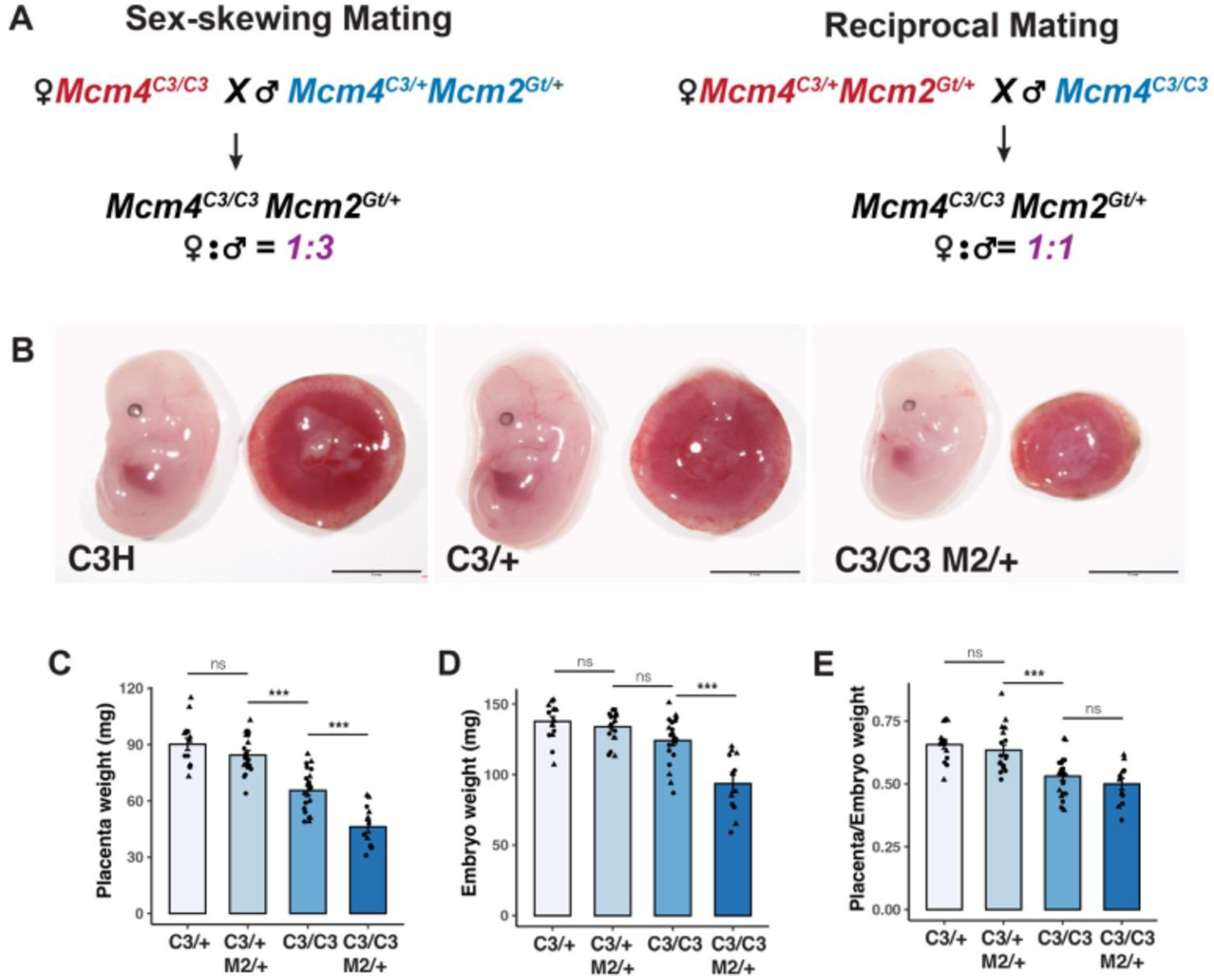
Reduced placental and embryonic weight in embryos with genomic instability. (A) Mating scheme for sex-skewing and reciprocal crosses to generate high GIN mutants. (B) Representative images of embryos with indicated genotypes. Scale bar: 5mm. (C-E) Placental weight, embryonic weight, and placental-to-embryonic weight ratio at E13.5 from sex-skewing matings. C3/+: *Mcm4^C3/+^*; C3/C3: *Mcm4^C3/C3^*; C3/+ M2/+: *Mcm4^C3/+^ Mcm2^Gt/+^*; C3/C3 M2/+: *Mcm4^C3/C3^ Mcm2^Gt/+^*. p-values were calculated with one-way ANOVA with Tukey’s HSD test. ns: not significant. *: p<0.05; **: p<0.01; ***: p<0.001. Error bars: SEM. Each data point represents a single placenta or embryo; females (circles), males (triangles).

To further investigate the cause of female-biased embryonic lethality, we assessed embryonic and placental growth during mid-gestation in both sex-skewing and reciprocal matings (Fig. 1A). At E13.5, *Mcm4^C3/+^* embryos and placentae from the sex-skewing mating were grossly indistinguishable from WT (produced from WTxWT matings) (Fig. 1B). However, *Mcm4^C3/C3^ Mcm2^Gt/+^* placentae and embryos from the sex-skewing mating were smaller than *Mcm4^C3/+^* littermates and *Mcm4^+/+^*controls (Fig. 1B). Both embryonic and placental weights were significantly reduced in *Mcm4^C3/C3^ Mcm2^Gt/+^* genotypes from both sex-skewing and reciprocal matings (Fig. 1C-E; Supplemental Fig. 1A-C), but the reductions in females were more severe in sex-skewing matings than in reciprocal matings. Males of the same genotype didn’t show parent-of-origin dependent placental weight reduction (Supplementary Fig. 1D). In addition, the placenta-to-embryo weight ratio was significantly reduced in *Mcm4^C3/C3^* and *Mcm4^C3/C3^ Mcm2^Gt/+^* genotypes from both sex-skewing and reciprocal matings at E13.5, suggesting that the growth in placenta was significantly hindered in high GIN genotypes (Fig. 1E; Supplemental Fig. 1C).

To investigate whether the effect of GIN on placental development is unique to the *Chaos3* model, we collected placentae and embryos from *Fancm^+/-^*intercrosses during mid-gestation and measured their weights. FANCM is involved in DNA replication fork repair. Similar to *Chaos3* mice, FANCM deficiency also presents with increased GIN and female-biased lethality ^17,23^. At E13.5, both placental and embryonic weights of *Fancm^-/-^* animals were significantly reduced compared to control littermates (Supplemental Fig. 2A, B). These data suggested that the developing trophoblast lineage is particularly sensitive to persistent GIN.

### Semi-lethal mutants exhibit placental developmental defects preferentially affecting the junctional zone (JZ)

To determine whether the reduction in placental weight in semi-lethal genotypes was related to any underlying structural defects, we carried out histological analysis of placentae at E13.5. Periodic Acid Schiff (PAS) staining showed significantly reduced JZ and labyrinth zone (LZ) areas in semi-lethal genotype placentae compared to control littermates from both sex-skewing and reciprocal matings (Fig. 2A-C, Supplementary Fig. 3A-C). Within the JZ, PAS-positive glycogen trophoblast cells (GlyTs) were almost absent in *Mcm4^C3/C3^ Mcm2^Gt/+^* placentae from sex-skewing matings (Fig. 2A). Additionally, spongiotrophoblasts (SpTs) and parietal Trophoblast Giant Cells (p-TGCs, which have distinctly large nuclei compared to diploid cells) within the central regions of the maternal-fetal interface were severely depleted (Fig. 2A). On the other hand, some SpTs were present in the *Mcm4^C3/C3^ Mcm2^Gt/+^* placentae from reciprocal matings (Supplementary Fig. 3A). Additionally, we found that the reduction in the JZ area was more prominent in females with the semi-lethal genotype compared to males from sex-skewing matings (Supplementary Fig. 3E-G). In contrast, no sex differences in JZ areas were observed in placentae from reciprocal matings (Supplementary Fig. 3E). The JZ:total placental area (JZ+LZ) ratio was significantly reduced in semi-lethal genotypes (Fig. 2D) with a more prominent reduction in females from sex-skewing matings (Supplementary Fig. 3G). Female, but not male, *Fancm^-/-^* placentae also had a reduced JZ, but the LZ appeared normal (Supplementary Fig. 2C, D).

**Figure 2.**
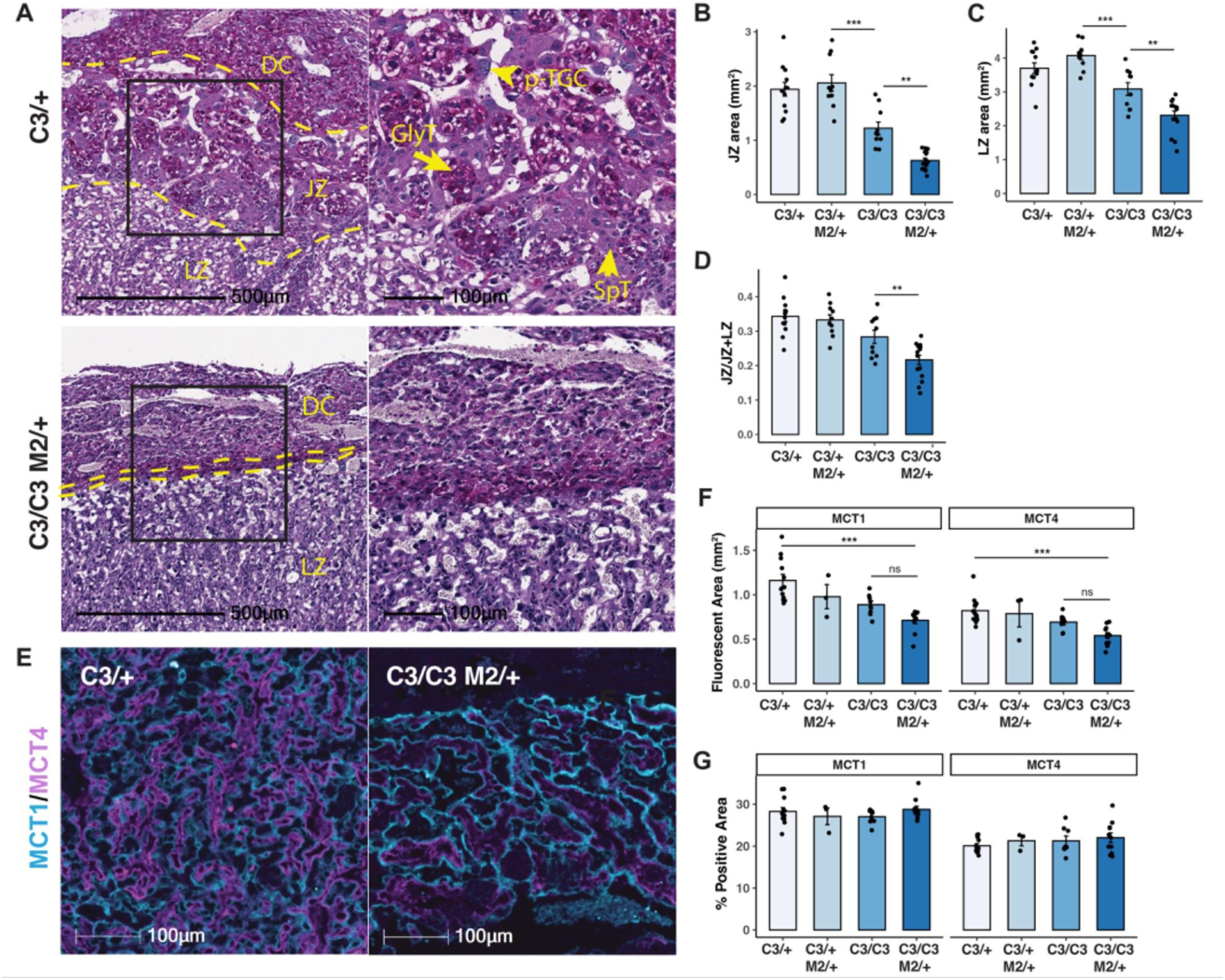
Placentae with replication stress have smaller junctional and labyrinth zones. (A) Periodic Acid-Schiff staining of E13.5 placental sections from indicated genotypes. The middle and left panels show magnified views of the boxed regions in the right panel. (B-D) Quantification of junctional zone (JZ) and labyrinth zone (LZ) areas and JZ proportion in placentae from sex-skewing matings at E13.5. (E) Representative MCT1 and MCT4 staining of E13.5 placental sections. (F) Quantification of MCT1 and MCT4 fluorescent areas, and (G) the percentage of positive areas in placental sections of indicated genotypes.C3/+: *Mcm4^C3/+^*; C3/C3: *Mcm4^C3/C3^*; C3/+ M2/+: *Mcm4^C3/+^ Mcm2^Gt/+^*; C3/C3 M2/+: *Mcm4^C3/C3^ Mcm2^Gt/+^*. DC: decidua; JZ: junctional zone; LZ: labyrinth zone; p-TGC: parietal trophoblast giant cells; SpT: spongiotrophoblast; SynT: syncytiotrophoblast; GlyT: glycogen trophoblast. p-values were calculated with one-way ANOVA with Tukey’s HSD test. ns: not significant. *: p<0.05; **: p<0.01; ***: p<0.001. Error bar: SEM. Each data point represents the average of at least three sections from the same placenta.

To determine whether the diminished LZ was due to a decreased number of trophoblasts, we immunostained E13.5 placentae with MCT1 and MCT4 that are markers of Syncytiotrophoblasts I and II (SynTI and SynTII) respectively. The overall MCT1- and MCT4-positive areas were reduced in *Mcm4^C3/C3^ Mcm2^Gt/+^* genotypes compared to control littermates (Fig. 2E), in agreement with the observed reduction in LZ size. However, the proportions of MCT1 and MCT4 positive areas within the LZ in *Mcm4^C3/C3^ Mcm2^Gt/+^* genotypes were not different from the control littermates (Fig. 2F, G).

We next performed bulk RNA-Seq of E13.5 placental samples as a means to corroborate the histological observations, focussing on the expression of genes characteristic of various trophoblast cell types as revealed by single-cell RNA-Seq studies ^24^. Consistent with our histological observations, expression of key JZ trophoblast cell markers (e.g., P-TGC: *Prl3b1*, *Prl2c2*; SpT: *Prl8a8*, *Tpbpa*; GlyT: *Gjb3*, *Prl6a1*; SpA-TGC: *Plac8*, *Rgs5*) in *Mcm4^C3/C3^ Mcm2^Gt/+^* placentae were reduced vs WT, and particularly in female placentae (Supplementary Fig. 3H). In contrast, expression of LZ trophoblast markers (e.g., S-TGC: *Ctsq*, *Hsd17bq*; SynTI: *Glis1*, *Stra6*; SynTII: *Gcm1*, *Gcgr*) were similar to the WT and no sex differences were observed (Supplementary Fig. 4A). Furthermore, the percentage of proliferating cells in the JZ, but not in the LZ, was significantly reduced in *Mcm4^C3/C3^ Mcm2^Gt/+^* placentae as indicated by Ki67 immunostaining, and was negatively correlated with the level of GIN in litters from the sex-skewing mating (Fig. 3A, B). Collectively, these data suggest that the JZ trophoblast cells are particularly susceptible to chronic RS mediated by MCM deficiency.

**Figure 3.**
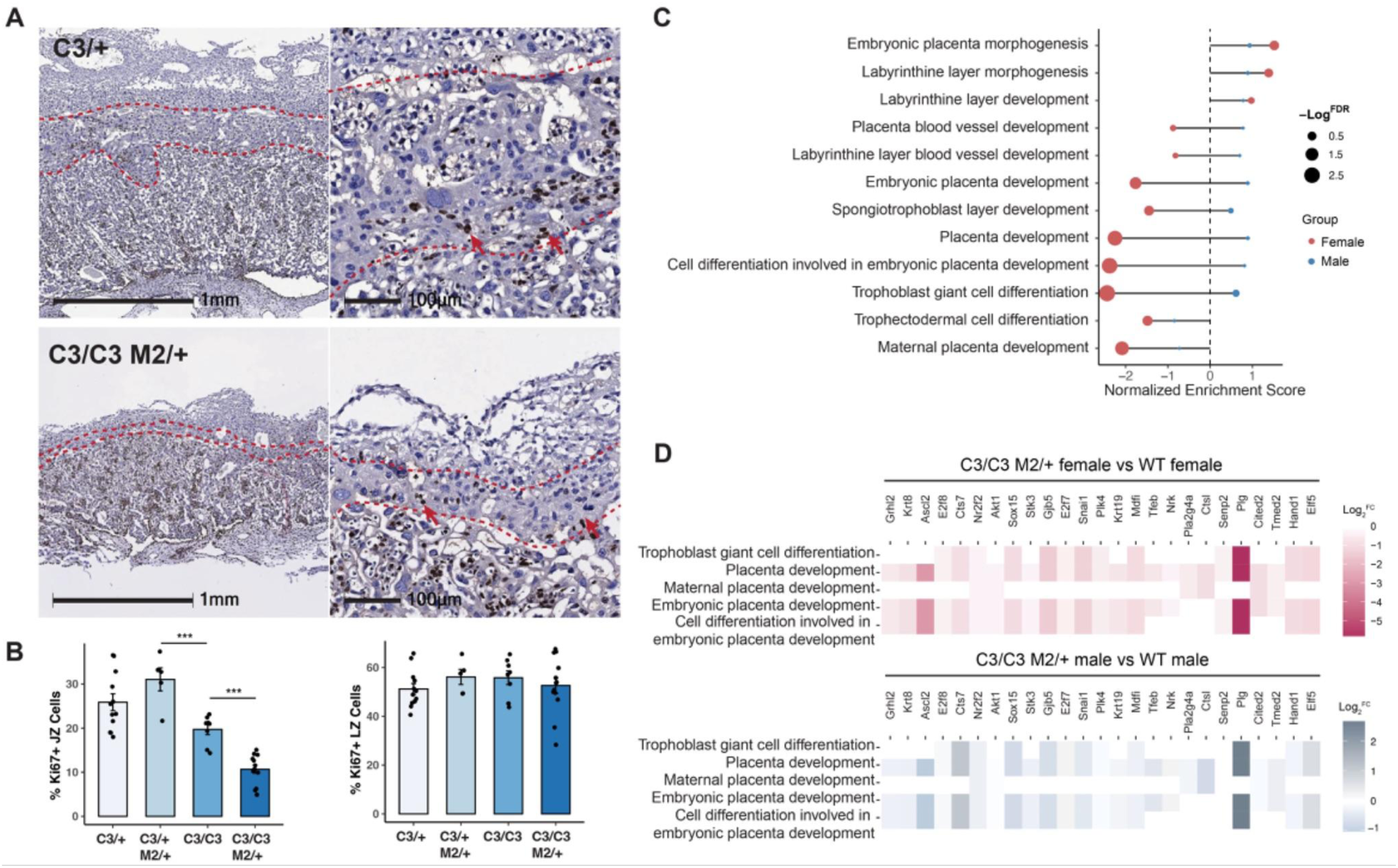
Impact of GIN on LZ is less than JZ. (A) Representative Ki67 immunohistochemistry of E13.5 placental sections from indicated genotypes. (B) Quantification of Ki67-positive cells in the JZ and LZ from sex-skewing matings. Error bar: SEM. Each data point represents the average of at least three sections from the same placenta. (C) Gene Ontology (GO) enrichment analysis of placenta development-related terms. (D) Log-transformed fold change of shared placental development genes in females and males. C3/+: *Mcm4^C3/+^*; C3/C3: *Mcm4^C3/C3^*; C3/+ M2/+: *Mcm4^C3/+^ Mcm2^Gt/+^*; C3/C3 M2/+: *Mcm4^C3/C3^ Mcm2^Gt/+^*. p-values were calculated with one-way ANOVA with Tukey’s HSD test. ns: not significant. *: p<0.05; **: p<0.01; ***: p<0.001.

Additional gene set enrichment analysis (GSEA) of the bulk RNA-Seq data reflected disruption of the trophoblast compartment. Gene Ontology (GO) biological processes related to placental development in female, but not male, *Mcm4^C3/C3^ Mcm2^Gt/+^* placentae were significantly underrepresented, such as trophoblast giant cell differentiation and spongiotrophoblast layer development (Fig. 3C). Consistent with the histological assessment, expression of genes involved in these underrepresented biological processes were significantly lower in female *Mcm4^C3/C3^ Mcm2^Gt/+^* placentae compared to WT (Fig. 3D). These differences were less pronounced in male mutants (Fig. 3D). Some of the shared genes in these pathways are predominantly expressed in the JZ trophoblast, such as *Ascl2 ^25^, Gjb5 ^26^*, and *Hand1 ^27^*, the deficiency of which result in poor JZ development ^26,28,29^.

Despite the disrupted LZ in *Mcm4^C3/C3^ Mcm2^Gt/+^* placentae, pathways related to angiogenesis were overrepresented compared to WT (Supplementary Fig. 4B). Interestingly, this was more prominent in female mutants (Supplementary Fig. 4C). For instance, *Vegfa* and *Kdr* transcripts were significantly elevated in mutant female placentae compared to WT or mutant males (Supplementary Fig. 4C). Given the dramatic LZ reduction in the mutants, the overrepresentation of angiogenic pathways might suggest an underlying tissue hypoxia due to defective branching morphogenesis, as seen in the *Mcm4^C3/C3^ Mcm2^Gt/+^* placentae (Fig. 2E). Alternatively, delayed development of the LZ might result in transcriptomic profiles reflecting earlier stages of labyrinth development. In summary, these data suggest that persistent RS exerts lineage-specific, sexually dimorphic impacts on placental development.

### GIN impairs trophoblast stem cell (TSC) establishment and maintenance

In most cell types, high levels of DNA damage result in the activation of DNA damage checkpoints causing cell cycle arrest or apoptosis ^4^. At E11.5, we found no evidence of elevated apoptosis (Supplementary Fig. 5A) in the mutant placentae. However, JZ cells were almost absent at this stage, suggesting that defects in the spongiotrophoblast layer occurred earlier during development. At E9.5, TPBPA, a marker of JZ trophoblast, was expressed in a smaller region of *Mcm4^C3/C3^ Mcm2^Gt/+^* placentae compared to control littermates, and few apoptotic cells were observed in the spongiotrophoblast layer (Supplementary Fig 5B, C). These data suggest that defects in *Mcm4^C3/C3^ Mcm2^Gt/+^*placentae likely occurred early, either from impaired differentiation ability of trophoblast progenitors, or defects in the initial maintenance of the TSC pool.

To test these hypotheses, we performed experiments to derive mutant TSC lines that could be used for growth and differentiation studies. Blastocysts obtained from intercrosses of *Mcm4^C3/+^* mice were cultured in conventional TSC derivation media containing FGF4 and heparin on a mouse embryonic fibroblast (MEF) feeder layer ^30^. The proportion of *Mcm4^C3/C3^*TSCs obtained was much lower than the expected Mendelian ratio (Supplementary Table 2), and these lines were difficult to maintain. In an attempt to improve derivation and/or growth efficiency, we next derived TSCs on Matrigel in a chemically defined medium containing growth factors and inhibitors ^31,32^ (Supplementary Fig. 6A). Again, *Mcm4^C3/C3^* lines were established at sub-Mendelian ratios (Supplementary Table 2), but they proliferated more stably under chemically defined conditions. Attempts to derive *Mcm4^C3/C3^ Mcm2^Gt/+^* TSC lines from crosses of *Mcm4^C3/C3^* females to *Mcm4^C3/C3^ Mcm2^Gt/+^* males were only successful using the defined condition with Matrigel. Thus, high RS is a major impediment to TSC growth or maintenance at very early stages, at least *in vitro*.

To evaluate the differentiation capacity of mutant TSCs, we initiated spontaneous differentiation by removing all the growth factors and inhibitors from the TSC culture medium. Starting on day 7 (D7) of differentiation, TGCs (PL-1) and SpTs, GlyT (TPBPA) markers can be seen with subsequent loss of the TSC marker EOMES (Supplementary Fig. 6B). However, by day 10 (D10), differentiated *Mcm4^C3/C3^* TSCs exhibited reduced levels of the TGC marker PL-1 compared to WT (Supplementary Fig. 6B).

To assess the impact of RS on TSC proliferation, we performed EdU pulse-labeling. Both *Mcm4^C3/C3^* and *Mcm4^C3/C3^ Mcm2^Gt/+^* TSCs displayed significantly fewer EdU^+^EOMES^+^ cells than WT, indicating a diminished proliferative capacity (Fig. 4C, D). In parallel, cleaved-Caspase 3 staining revealed a substantial increase of apoptotic cells in both *Mcm4^C3/C3^* and *Mcm4^C3/C3^ Mcm2^Gt/+^* TSCs (Fig. 4E, F), coupled with a reduced percentage of EOMES^+^ cells compared to WT TSCs (Fig. 4G, H), suggesting a loss of stemness in TSCs with high GIN.

**Figure 4.**
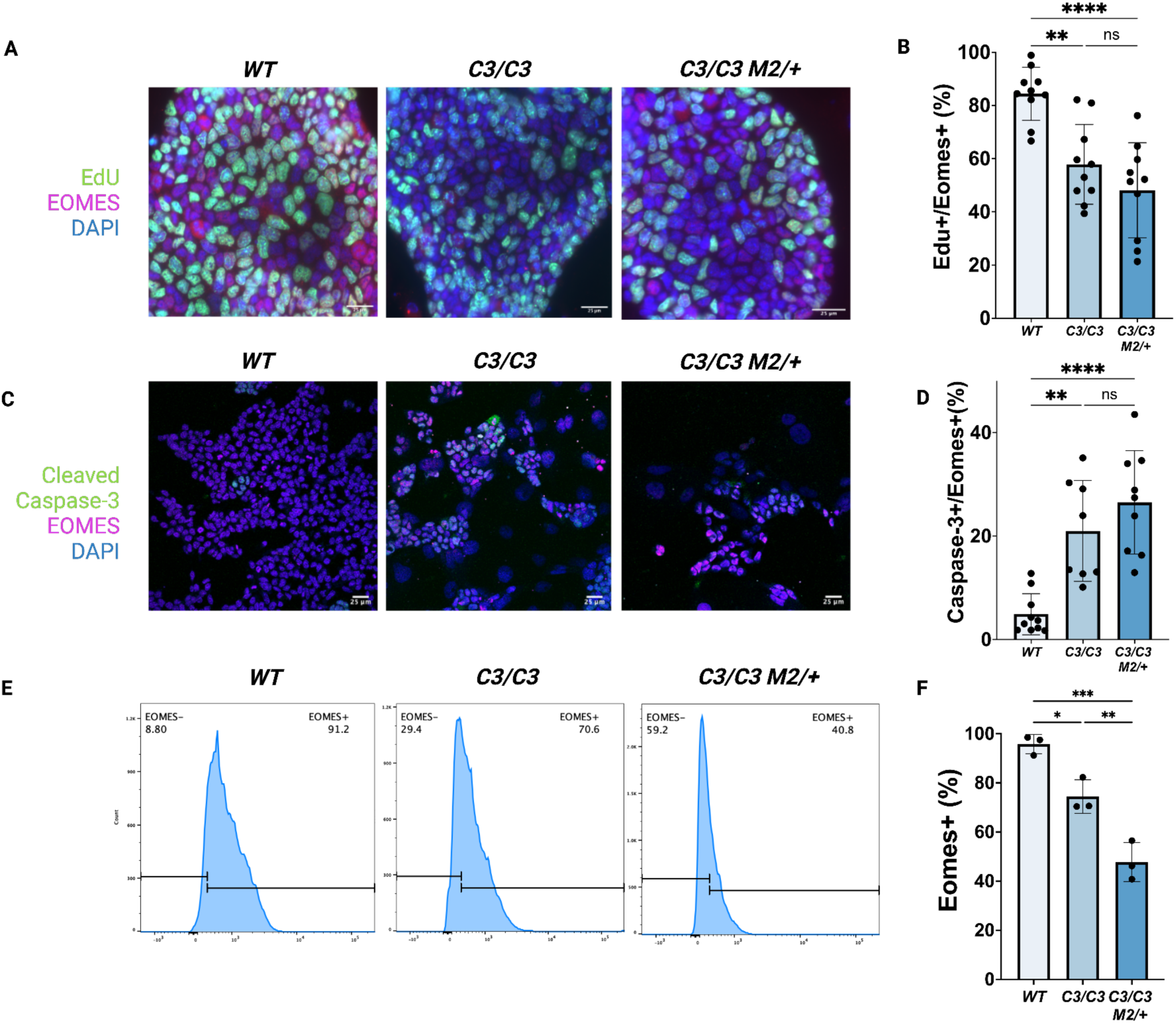
Trophoblast stem cells (TSCs) with increased genomic instability exhibit abnormal phenotypes. (A) EdU pulse labeling of TSCs with immunofluorescence staining for the EOMES. Scale bar: 25 µm. (B) Quantification of EOMES and EdU-double positive cells. (C) Immunofluorescence staining of TSCs for the Cleaved Caspase-3 and EOMES. (D) Quantification of EOMES and Cleaved Caspase-3-double positive cells. (E) Flow cytometry analysis of EOMES-positive TSCs. (F) Quantification of EOMES-positive cells. C3/+: *Mcm4^C3/+^*; C3/C3: *Mcm4^C3/C3^*; C3/C3 M2/+: *Mcm4^C3/C3^ Mcm2^Gt/+^*. The dots in (B), (D) and (F) represent individual biological replicates from independent experiments. p-values were determined by one-way ANOVA with Tukey’s HSD test. Error bars represent mean ± SEM. *p < 0.05, **p < 0.01, ***p < 0.001, ****p<0.0001; ns: not significant.

In contrast to the deficiencies in mutant TSC maintenance, both *Mcm4^C3/C3^* and *Mcm4^C3/C3^ Mcm2^Gt/+^* mouse embryonic stem cell (ESC) lines were readily established from blastocysts and successfully maintained; we observed no differences in the expression of naive pluripotency marker KLF4 among WT, *Mcm4^C3/C3^* and *Mcm4^C3/C3^ Mcm2^Gt/+^* ESCs (Supplementary Fig. 7A, B). Proliferation rate, assessed by the proportions of EdU+KLF4+ cells, were also comparable between GIN genotypes ESCs and WT ESCs (Supplementary Fig. 7C, D). Consistent with the previous findings ^33^, our data suggest that MCM deficiency does not affect ESCs pluripotency or self-renewal. In summary, we found that persistent RS impairs TSC, but not ESC, viability, proliferation and maintenance *in vitro*.

### RS-induced checkpoint activation and CDK1 inhibition drive premature differentiation in TSCs

To further elucidate the possible molecular mechanisms underlying the observed defects in TSCs with high GIN, we hypothesized that TSCs activate cell cycle checkpoints via the DNA damage response (DDR). As previously demonstrated for mutant placentae ^17^, both *Mcm4^C3/C3^* and *Mcm4^C3/C3^ Mcm2^Gt/+^* TSCs contained substantially more γH2AX (a marker of strand breaks, primarily DSBs) than WT cells (Fig. 5A). In contrast, ESCs bearing the GIN genotypes showed a similar percentage of γH2AX positive cells and the average intensity of γH2AX remained the same across WT, *Mcm4^C3/C3^* and *Mcm4^C3/C3^ Mcm2^Gt/+^* ESCs (Supplementary Fig. 7E-G). This suggests that ESCs are less sensitive than TSCs to DNA damage induced by RS, or do not have a robust DDR (as indicated by H2AX S139 phosphorylation) as a consequence of MCM deficiency. Furthermore, cultures of TSCs with high levels of GIN were enriched for the G2/M population, suggesting the activation of checkpoint signaling pathways (Fig. 5C, D). In addition, both *Mcm4^C3/C3^* and *Mcm4^C3/C3^ Mcm2^Gt/+^* TSCs displayed a significant increase in the >4C population relative to WT (Fig. 5C, E), suggestive of increased endoreduplication, a process in which repeated rounds of DNA replication occur without mitosis, and which is a remarkable characteristic of mouse TGCs ^12^.

**Figure 5.**
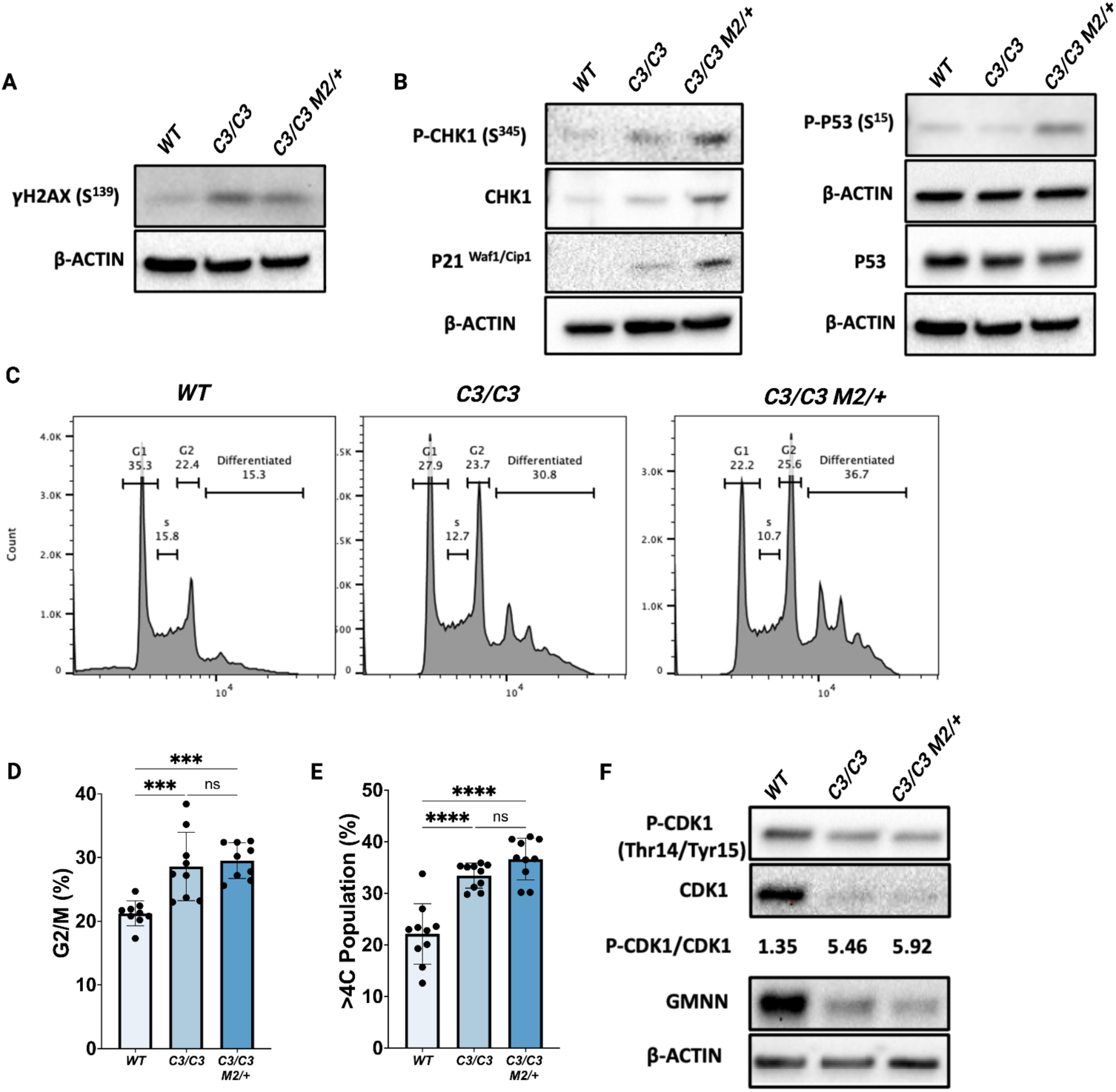
DNA damage responses and G2/M checkpoint activation drive premature TSC differentiation. (A) Western blot analysis of γH2AX (S139) levels in TSCs of the indicated genotypes. (B) Western blot analysis of DNA damage checkpoint proteins in TSCs. (C) Flow cytometry analysis of DNA content in TSCs. The >4C populations are labeled as “differentiated.” (D-E). Quantification of (D) DNA content and (E) differentiated (>4C) cell populations across genotypes. (F) Western blot analysis of G2/M regulators in TSCs of the indicated genotypes. *C3/C3*: *Mcm4^C3/C3^*; *C3/C3 M2/+*: *Mcm4^C3/C3^ Mcm2^Gt/+^*.The dots in (D) and (E) represent individual biological replicates from independent experiments. p-values were determined by one-way ANOVA with Tukey’s HSD test. Error bars represent m ean ± SEM. *p < 0.05, **p < 0.01, ***p < 0.001, ****p<0.0001; ns: not significant.

Given the observed cell cycle disruption in TSCs, we investigated the signaling pathways involved in regulating the G2/M transition. Western blot analysis revealed elevated levels of phosphorylated CHK1 (S345), phosphorylated p53 (S15), and p21 in *Mcm4^C3/C3^* and *Mcm4^C3/C3^ Mcm2^Gt/+^* TSCs compared to WT (Fig. 5B). Furthermore, mutant TSCs exhibited a reduction in Cyclin-dependent kinase 1 (CDK1) levels, along with an increased Phospho-CDK1 (Thr14/Tyr15) to total CDK1 ratio, suggesting enhanced CDK1 inhibition (Fig. 5F). Inhibition of CDK1, which drives the G2/M transition and is a downstream target of p21 (CDKN1A), is required for cells to enter endoreduplication and thereby TGC differentiation ^9,34^. Collectively, these findings indicate that persistent RS in TSCs activates the CHK1-p53-p21 axis, leading to CDK1 inhibition and premature differentiation via endoreduplication. Normally, eukaryotic cells have a robust mechanism to ensure that the genome is replicated only once per cell cycle by preventing origin licensing after S phase initiation, and Geminin (GMNN) is a key factor in this process ^35, 36^. We observed reduced GMNN in *Mcm4^C3/C3^* and *Mcm4^C3/C3^ Mcm2^Gt/+^* TSCs (Fig. 5E), which may facilitate their endoreduplication and premature differentiation into TGCs.

### Decreasing MCM nuclear export factor MCM3 genetically rescues placental defects

*In vivo*, reducing MCM3 levels via heterozygosity paradoxically improved phenotypes associated with *Chaos3* mutants ^17,21,22^. In addition to a reduction in cancer occurrence, *Mcm3* heterozygosity doubled the percentage of viable *Mcm4^C3/C3^ Mcm2^Gt/+^* animals at birth ^21,22^. It was hypothesized that reducing MCM3, which is an MCM nuclear export factor in yeast ^37^, increased available MCMs for chromatin loading, thereby ameliorating RS ^21^. Reduced MCMs in mice have been associated with deficits in stem cell compartments ^38,39^, and increasing the availability of chromatin-bound MCMs have been shown to improve the reprogramming efficiency of mouse embryonic fibroblasts ^21^. Therefore, we reasoned that if the loss in cellularity in the JZ in MCM-deficient *Mcm4^C3/C3^ Mcm2^Gt/+^* placentae was indeed caused by defective TSC maintenance, then reducing MCM3 in the *Mcm4^C3/C3^ Mcm2^Gt/+^* background might partially rescue placenta phenotypes. Indeed, the placenta and embryo weights of E13.5 *Mcm4^C3/C3^ Mcm2^Gt/+^ Mcm3^Gt/+^* animals were significantly improved *vs. Mcm4^C3/C3^ Mcm2^Gt/+^* animals, rendering them similar to *Mcm4^C3/C3^* littermates (Fig. 6A-C). The JZ (but not LZ) area in *Mcm4^C3/C3^ Mcm2^Gt/+^ Mcm3^Gt/+^* placentae also increased to the level of *Mcm4^C3/C3^* littermates (Fig. 6D-H). These data suggest that the placental abnormalities were exacerbated by reduced MCMs, which is known to cause elevated GIN from insufficient numbers of dormant (“backup”) origins ^19,20,40^.

**Figure 6.**
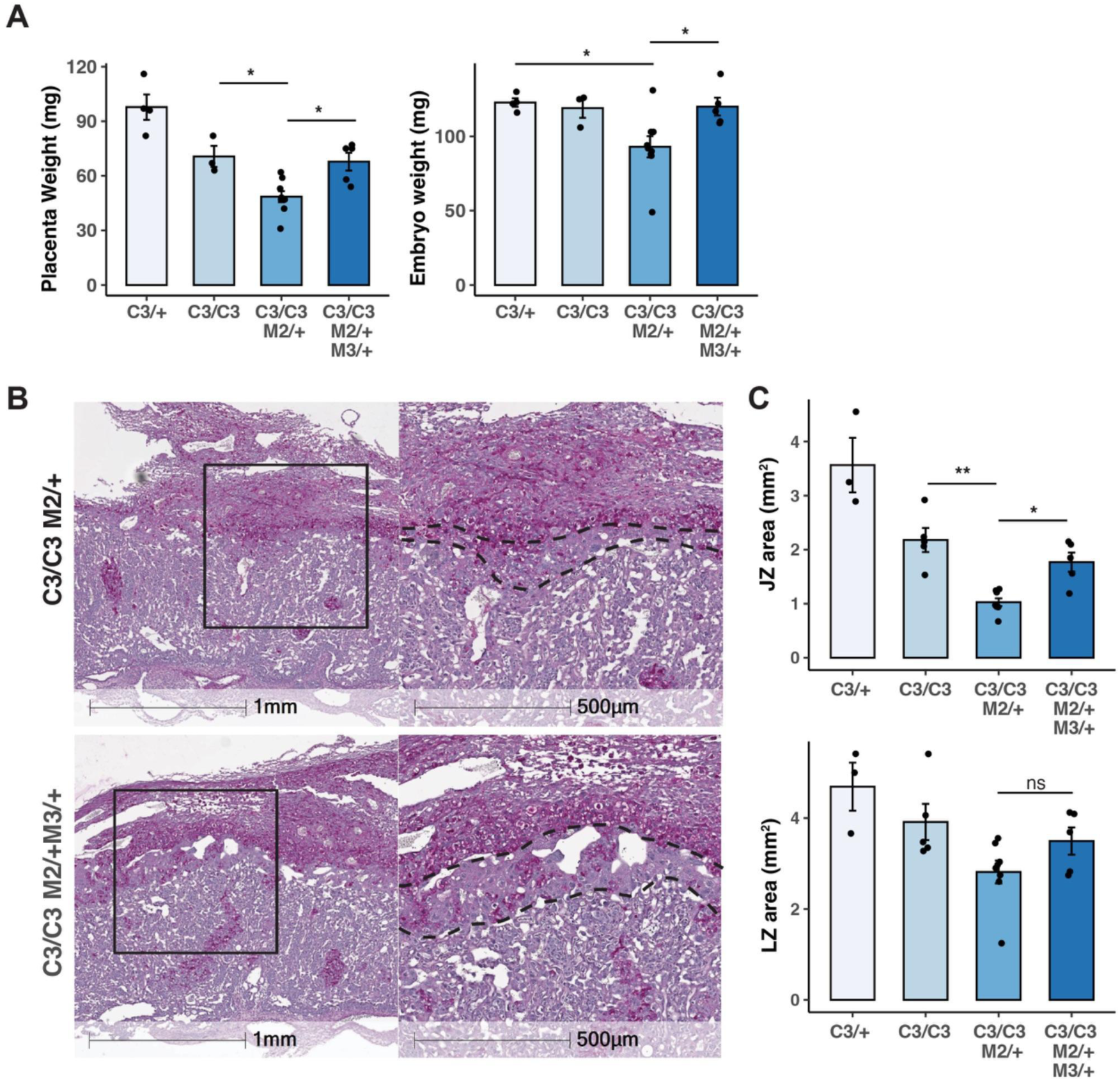
*Mcm3* heterozygosity rescues placental defects. (A) Plots of placental and embryonic weights. Each data point is one placenta or an embryo. (B) PAS staining of E13.5 placental sections. (C) Average sizes of JZ and LZ areas. Each data point represents the average measurement taken from at least three sections of the same placenta. C3/+: *Mcm4^C3/+^*; C3/C3: *Mcm4^C3/C3^*; C3/C3 M2/+: *Mcm4^C3/C3^Mcm2^Gt/+^*; C3/C3 M2/+ M3/+: *Mcm4^C3/C3^Mcm2^Gt/+^ Mcm3^Gt/+^*. *p-values* were determined by one-way ANOVA with Tukey’s HSD test. ns: not significant. *: p<0.05; **: p<0.01. Error bar: SEM.

## Discussion

Proper placentation is a prerequisite for embryo survival and a healthy pregnancy. Defects in the development of trophoblast cell types have been linked to pregnancy complications endangering both the mother and the fetus ^41,42^. Here, we show that GIN caused by either MCM or FANCM deficiencies impairs JZ development (Fig. 2A-D, Supplementary Fig.2). The labyrinth histology in GIN mutants exhibited reduced complexity, indicative of poor branching morphogenesis and/or underdeveloped blood spaces (Fig. 2E-G). However, the impact of MCMs deficiency seemed to be more severe in the JZ than the labyrinth as the percentage of proliferating cells in the LZ were not affected (Fig. 3A, B). The JZ cells act as hormonal and energy reservoirs for normal fetal development, and a smaller JZ is associated with intrauterine growth restriction (IUGR) ^42^. Indeed, semilethal genotype embryos were significantly smaller compared to control littermates at mid-gestation (Fig. 1C-H), and those that survived to birth also suffered from growth retardation ^22^.

Mutations decreasing MCM levels have been associated with reduced stem cell fitness ^39,43^, and our studies indicate that normal levels and biochemical functions of MCMs in mouse TSCs are crucial for proper placentation and embryogenesis. In mice, almost all JZ trophoblasts originate from cells in the ectoplacental cone, which arises from continued expansion of the extraembryonic ectoderm between E5.5 and E7.5 ^41^. Polar trophectoderm in the developing blastocyst and cells in the extraembryonic ectoderm are considered to be the stem progenitors of all trophoblast cells that contribute to the mature placenta ^41^. Under persistent RS, the percentage of proliferating cells in the EPC and ExE were significantly reduced at E7.5 (Supplementary Fig. 5D, E). TSC lines with high levels of GIN were derived inefficiently, and under conventional culture conditions, difficult to maintain, likely because such conditions do not adequately suppress differentiation signals ^30^. Growth factors and inhibitors such as Wnt inhibitor, ROCK inhibitor, and Activin A are often required to block differentiation pathways and promote self-renewal ^31,32^. Use of such conditions enabled us to derive *Mcm4^C3/C3^ Mcm2^Gt/+^* mutant TSCs, although they exhibited reduced proliferation potential and increased cell death (Fig. 4D-F). Additionally, the observation of increased cells with >4C DNA content in mutant TSC cultures is consistent with premature differentiation into TGCs. In contrast, ESCs bearing the same GIN genotypes did not exhibit reduced proliferation, poor stem cell maintenance or signs of DNA damage (Supplementary Fig. 7). This is probably because ESCs have characteristics rendering them relatively resistant to RS, namely high levels of licensed dormant origins, rapid origin licensing due to having a short G1 phase, and unique replication-coupled mechanisms to safeguard their genomes ^33,44,45^.

We propose that TSC defects in GIN genotypes were due to CDK1 inhibition resulting from activation of the CHK1-p53-p21 checkpoint pathway (Fig. 5). Inhibition of CDK1, a key regulator of the G2/M transition, is sufficient to drive endoreduplication, promoting TSCs to differentiate into TGCs ^9,34^. Consistently, we observed reduced CDK1 activity in TSCs with high levels of GIN (Fig. 5F). On the other hand, TSCs were able to differentiate into JZ cells, namely SpTs and TGCs when prompted to differentiate (Fig. 4C, TPBPA and PL1 immunoblots respectively). Taken together, we propose that due to poor TSC maintenance in semi-lethal genotypes, there was an insufficient pool of EPC cells that ultimately contributed to the poor cellularization of the JZ in the mature placenta. Consistent with the importance of robust MCM origin licensing for stem cell maintenance, the placental size and the JZ defects were partially rescued by breeding mice that genetically rescue viability, presumably by increasing chromatin bound MCMs as shown in mutant MEFs (Fig. 6) ^21,22^.

We were unable to rescue the viability of semilethal genotype embryos by deleting cytosolic nucleotide sensing pathways (Supplementary Table 1). Consistent with this, a recent study showed that endogenous cytosolic DNA/micronuclei cannot activate cGAS-STING in primary and cancer cell lines ^46^. This suggests that the nature of inflammation in the placenta may not be solely related to GIN *per se*, but possibly the underlying structural anomalies (Fig. 2). We showed previously that treating pregnant dams with ibuprofen abolished sex bias and the viability of both male and female *Mcm4^C3/C3^ Mcm2^Gt/+^* embryos was partially improved ^17^. However, ibuprofen treatment had very little impact on placental structure (not shown), suggesting that the action of ibuprofen is likely a downstream symptomatic relief of the impact resulting from the structural anomalies. Indeed, reduction in the JZ was more severe in the female semi-lethal genotype placentae than the males. Furthermore, genes predominantly expressed in the JZ cells were significantly underrepresented in the female placentae with semi-lethal genotypes (Fig. 3C, D). Although the exact underlying mechanism of such sexual dimorphism needs further investigation, maternal factors, such as the epigenetic information carried by the oocyte, might account for sex-specific and parent-of-origin dependent placental defects ^47^. Nevertheless, further experiments are needed to determine whether placental defects are the main cause of female biased embryonic semi-lethality in the *Chaos3* model.

MCMs are absolutely essential for DNA replication and the maintenance of genomic stability. While null alleles of any of the 6 MCMs cause pre-implantation lethality, hypomorphic alleles - such as the *Chaos3* allele of *Mcm4* - have enabled identification of tissue-specific sensitivities and impacts of compromised DNA replication ^17,39,43^. Death of *Mcm4^C3/C3^ Mcm2^Gt/+^* embryos are first detected after E9.5, a period in mouse development where embryonic lethality is typically associated with placental defects ^48^. The Mouse Genome Informatics (MGI) database lists 106 genes (out of ∼ 900) involved in genome maintenance whose deficiency causes embryonic lethality between E9.5-E14.5. Among these, only 26 of them were reported to have placentation defects, and the underlying molecular mechanism needs investigation (Supplementary Table 3). Furthermore, this is likely an underestimate as 70% of mouse knockout alleles that are embryonic lethal around this stage were reported to have abnormal placentation ^48^. Taken together, we propose a modification to the conventional belief that trophoblast cells are resistant to GIN, instead genome maintenance mechanisms are likely at play at certain stages of placenta development in a lineage dependent manner. Whether RS compromises human placentation needs exploration.

## Material and Methods

### Animal husbandry

Mouse studies in this work were approved by Cornell University’s Institutional Animal Care and Use Committee under protocol number 2004-0038. All husbandry and mating experiments are conducted in the same animal facility and room, and under the same environmental conditions. *Tmem173 ^Gt/Gt^* ^49^ (JAX stock #017537) and *Myd88 ^-/-^* ^50^ (JAX stock #009088) were purchased from the Jackson laboratory. *Ddx58 ^-/-^* line was generated by the Cornell Transgenic Core Facility using sgRNA 5’-GATATCATTTGGATCAACTG-3’. For timed matings, female (8-12 weeks old) and male (8-16 weeks old) mice were placed in mating cages one day prior to checking for vaginal plugs. Presence of vaginal plugs was designated as E0.5. Genotyping of tissue biopsies was performed by Transnetyx (Memphis, TN, USA) or as previously published ^22^ using primers listed in Supplementary Table 4.

### Histology

Placental samples were isolated from E13.5 embryos and cut in half at midline. Half of a placenta was snap frozen in liquid nitrogen for RNA extraction, while the other half was fixed overnight in 4% paraformaldehyde in 1X PBS at 4⁰C, dehydrated in 70% ethanol and embedded in paraffin blocks. Embedded samples were sectioned at 6µm thickness, deparaffinized in xylene, rehydrated in serial dilutions of ethanol solutions and finally in deionized water.

For Periodic Acid Schiff staining, rehydrated placental sections were incubated with 0.5% Periodic Acid for 5 minutes followed by 15 minutes incubation with Schiff’s reagent (PAS staining kit, Sigma-Aldrich, Cat#: 1.01646). Sections were dipped into Hematoxylin Gill II (Leica Biosystems) for 20 seconds followed by 5 minutes in tap water. Slides were scanned and imaged on a Leica Aperio CS2 scanner. For measuring the junctional and labyrinth zone areas, at least six consecutive placental sections from the midline were stained, and the area of every other section was measured using FiJi/ImageJ software, and the average area was presented for each placenta.

For immunohistochemistry, deparaffinized and rehydrated sections were incubated with 3% hydrogen peroxide to quench endogenous peroxidase, then permeabilized with 0.1% Triton-X100 in 1X PBS at room temperature for 10 minutes, blocked for one hour with 3% goat serum in 1X PBS at room temperature. Incubation with primary antibodies was carried out overnight at 4⁰C. Rabbit primary antibodies were detected with SignalStain Boost IHC detection reagent (CST Cat#: 8114). Sections were dipped into Hematoxylin Gill II (Leica Biosystems) for 20 seconds followed by 5 minutes in tap water. Slides were scanned using Leica Aperio CS2 scanner. Ki67 positive cells were quantified using QuPath (https://qupath.github.io/). All antibodies used in this study are listed in Supplementary Table 5.

### TSC derivation and culture

TSCs were derived from preimplantation E3.5 blastocysts. For the defined culture condition, individual E3.5 blastocysts were placed in 48-well plates containing a mouse embryonic fibroblast (MEF) feeder layer and cultured in the defined TS-stem medium (see below). By day 4, proliferating blastocyst outgrowths were observed. Between days 5 and 8 post-plating, these outgrowths were dissociated into single cells using Trypsin-EDTA. Initial TSC colonies appeared within four days after disaggregation. The feeder layers were kept during derivation, but were not used when growing established TSCs. The defined TS-stem medium (500 mL) was composed of 240 mL Neurobasal (Thermo Fisher; 21103049), 240 mL DMEM/F12 Ham’s 1:1 (Thermo Fisher; 11320033), 2.5 mL N2 supplement 100X (Gibco; 17502048), 5 mL B-27 supplement 50X (Gibco; 17504044), 5 mL KnockOut serum replacement (Gibco; 10828028), 2.5 mL penicillin-streptomycin (Thermo Fisher; 15140122), 2.5 mL GlutaMAX Supplement (Gibco; 35050061), 30% bovine serum albumin (Tribioscience; TBS8031, 0.05% final concentration), 1.5 × 10⁻⁴ M 1-thioglycerol (Sigma-Aldrich; 88640), 25–50 ng/mL recombinant mouse basic FGF (Peprotech; 450-33-50UG), 20 ng/mL Human/Mouse/Rat Activin A Recombinant Protein (Peprotech; 120-14E-100UG), 10 μM XAV939 (Selleckchem; S1180), and 5 μM Y27632 (Stemcell; 72304). The medium was refreshed every two days. For feeder-free culture, TSCs were plated on 0.2% Matrigel-coated dishes and incubated at 37 °C for at least 1.5 hours or overnight before further culture.

For differentiation, TSCs were cultured in RPMI 1640 (Gibco; 11875085), 50 μg/mL penicillin-streptomycin (Gibco; 15140122), 2 mM L-glutamine (Gibco; 35050061), 1 mM sodium pyruvate (Gibco; 11360070), 100 μM β-mercaptoethanol (Sigma; M752), and 20% fetal bovine serum (Cytiva HyClone; SH30071.03HI).

### ESC derivation and culture

ESCs were derived as previously described ^51^. Briefly, E3.5 blastocysts were seeded into individual wells of the 48-well plates containing MEF feeder layers. Between day 7-10, the blastocyst outgrowths were mechanically dissociated and seeded onto a new feeder plate. For regular maintenance of ESCs, the following ESCs media was used: DMEM (Corning, 10-017-CM) supplemented with 15% fetal bovine serum (Cytiva HyClone; SH30071.03HI), 1% penicillin-streptomycin (Gibco; 15140122), 1% sodium pyruvate (Gibco; 11360070), 1% MEM NEAA (Gibco, 11140050), 100 μM β-mercaptoethanol (Sigma; M752), 3 μM CHIR99021 (Tocris, 4423), 1 μM PD0325901, 10^3^ IU LIF (Peprotech, 250-02-25UG). For the initial establishment of the ESCs lines, the fetal bovine serum in ESCs media was replaced with the same percentage of KnockOut serum replacement (Gibco; 10828028).

### Immunofluorescence of cultured cells

Cells were grown on in-chamber coverslips (iBidi µ-Slide 8 Well high; 80806) for 48 hours at 37℃ with 5% CO_2_, fixed in 4% paraformaldehyde for 15 min at room temperature, permeabilized with 0.1% Triton X-100 in 1X PBS for 15 min at room temperature, and blocked with 5% goat serum for one hour at room temperature. The cells were incubated with the primary antibodies diluted in 0.1% BSA in 1X PBS overnight at 4⁰C, followed by the secondary antibody at room temperature for one hour. Nuclei were stained with DAPI diluted in 0.1% BSA in 1X PBS. Slides were visualized using a Zeiss LSM 710 Confocal Microscope. Primary and secondary antibodies are listed in Supplementary Table 5.

### Western blotting

TSCs were trypsinized then washed twice with ice-cold 1× PBS before lysis in RIPA buffer (Thermo Fisher; 89900) supplemented with cOmplete™ Mini Protease Inhibitor Cocktail (Roche; 11836170001) and PhosSTOP™ Phosphatase Inhibitor Cocktail (Roche; 4906845001). TSC samples were sonicated three times at 25% amplitude for 10 seconds each (Thermo Fisher) and centrifuged at 14,000 × g for 15 minutes at 4°C. The supernatant was collected, proteins were denatured at 95°C for 5 minutes in sample buffer, then run on Bio-Rad 4–20% Mini-PROTEAN® Precast Gels (BioRad, Cat#: 4561094) at 100V for 75 minutes, then transferred onto PVDF membranes (Millipore; IPVH00010) on ice at 90V for 90 mins. Immunoblotting was performed using primary and secondary antibodies listed in Supplementary Table 5. Signal detection was carried out using enhanced chemiluminescence (Pierce; 32109) and visualized using a Bio-Rad Gel Doc XR Imaging System and analyzed with Image Lab (BioRad) software.

### Analysis of DNA content and cell cycle distribution

TSCs and differentiated trophoblast cells were harvested by trypsinization, washed twice with cold PBS, and fixed in 70% ethanol at −20°C for a minimum of two hours to preserve cellular DNA content. Fixed cells were then resuspended in 500 µL of propidium iodide (PI) staining solution, consisting of 0.1% Triton X-100, 1 µg/mL PI (Invitrogen; P3566), and 100 µg/mL RNase A (Thermo Fisher; EN0531) in 1× PBS. Following incubation at room temperature for 30 minutes in the dark, samples (100,000 events per sample) were analyzed on a BD FACSymphony™ A3 Cell Analyzer. Cell cycle distribution was determined using FlowJo software.

### EdU pulse labeling

For labeling TSCs, cells were treated with 10 µM EdU (5-ethynyl-2’-deoxyuridine) for 2 hours, and ESCs were treated for 30 minutes. The cells were then fixed in 4% paraformaldehyde for 15 minutes at room temperature, permeabilized with 0.1% Triton X-100 in PBS for 15 minutes, and stained using the Click-iT EdU Alexa Fluor 488 Imaging Kit (Thermo Fisher Scientific, C10337). Antibody labeling of cells was carried out after EdU detection. Nuclei were counterstained with DAPI to visualize total cell numbers. For TSCs, EdU and antibody labeled cells on the slides were imaged on Zeiss Axio Imager epifluorescence microscope. For ESCs, cells were analyzed on a BD FACSymphony™ A3 Cell Analyzer. The proportion of EdU-positive TSCs was quantified using FIJI/ImageJ software. The proportion of EdU-positive ESCs were analyzed using FlowJo software.

### RNA-seq and data analysis

Total RNA was isolated from WT and *Mcm4^C3/C3^ Mcm2^Gt/+^* placentae at E13.5 using Zymo Quick-RNA mini-prep Kit (Zymo Research, Cat#: R1054). RNA sample quality was confirmed by spectrophotometry (Nanodrop) to determine concentration and chemical purity (A260/230 and A260/280 ratios) and with a Fragment Analyzer (Agilent) to determine RNA integrity. PolyA+ RNA was isolated with the NEBNext Poly(A) mRNA Magnetic Isolation Module (New England Biolabs). UDI-barcoded RNAseq libraries were generated with the NEBNext Ultra II Directional RNA Library Prep Kit (New England Biolabs). Each library was quantified with a Qubit (dsDNA HS kit; Thermo Fisher) and the size distribution was determined with a Fragment Analyzer (Agilent) prior to pooling. Libraries were sequenced on an Illumina NovaSeq6000 (2×150 PE reads). At least 20 million 150bp paired-end reads were generated per library. For analysis, reads were trimmed for low quality and adaptor sequences with TrimGalore v0.6.0, a wrapper for cutadapt and FastQC using parameters: -j 1 -e 0.1 --nextseq-trim=20 -O 1 -a AGATCGGAAGAGC --length 50 -- fastqc. Unwanted reads were removed with STAR v 2.7.0e using parameters: -- outReadsUnmapped Fastx and final reads were mapped to the mouse reference genome Ensembl GRCm38 using STAR v2.7.0e. SARTools and DESeq2 v1.26.0 were used to generate normalized counts and statistical analysis of differential gene expression. Gene set enrichment analyses were performed using iDEP 2.0 (https://doi.org/10.1186/s12859-018-2486-6).

### Statistics

Statistical tests were carried out using one-way analysis of variance followed by Tukey’s post hoc test or student’s t-test using either R or Graphpad Prism. Graphs were generated using R or Biorender.

## Competing interest

The authors declare no competing or financial interests.

## Acknowledgements

We thank R. Munroe and C. Abratte of the Cornell Transgenic Core Facility for generating *Ddx58* knockout mice, and J. Grenier from Cornell’s Genomics Facility at the Cornell Institute of Biotechnology for the RNA-seq experiments. We also thank Cornell imaging and flow cytometry core. This work is supported by grant 5R01HD107910 from the National Institute of Child Health and Human Development to J.C.S and a Druckenmiller Postdoc Fellowship from the New York Stem Cell Foundation to M.M.

## Notes

### Competing Interest Statement

The authors have declared no competing interest.

### Summary of Updates

Supplementary Figure 7 is added to support the conclusion that the trophoblast stem cells are more sensitive to persistent replication stress induced by MCMs deficiency.

