## Supplemental files for "Chronic replication stress-mediated genomic instability disrupts placenta development in mice"

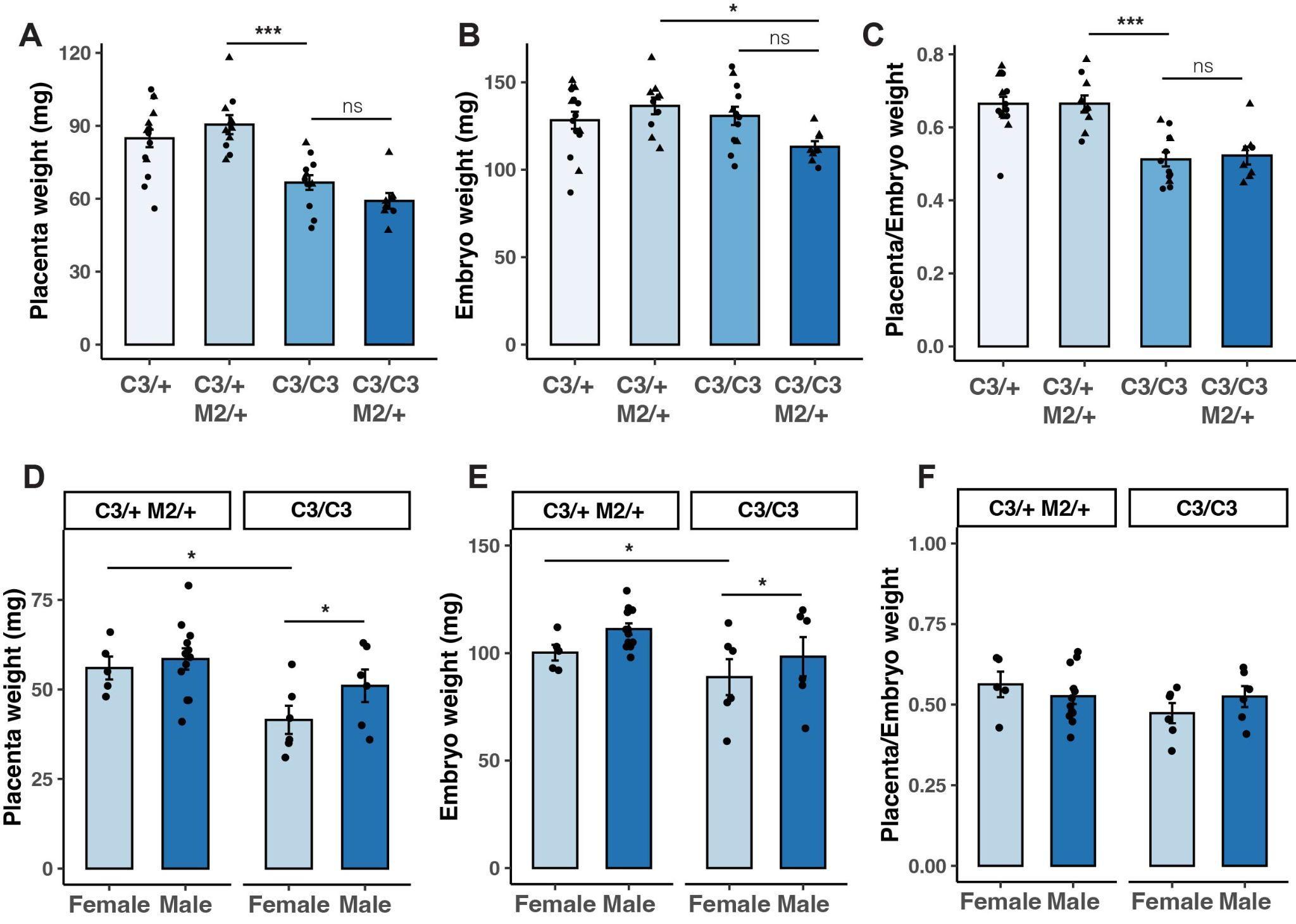

**Supplementary Figure 1. Embryos with genomic instability genotypes have reduced placental and embryonic weight.** (A-C) Placental and embryonic weight as well as placental-to-embryonic weight ratio of each genotype from the reciprocal mating at E13.5. Females (circles), males (triangles) (D-F). Comparison of placental and embryonic weight as well as placental-to-embryonic weight ratio of male and female *Mcm4^C3/C3^ Mcm2^Gt/+^* genotype from sex-skewing and reciprocal matings. Boxes indicating maternal genotype. C3/+: *Mcm4^C3/+^*; C3/C3: *Mcm4^C3/C3^*; C3/+ M2/+: *Mcm4^C3/+^ Mcm2^Gt/+^*; C3/C3 M2/+: *Mcm4^C3/C3^ Mcm2^Gt/+^*. p-values in (A-C) were calculated with one-way ANOVA followed by Tukey’s HSD test. p-values in (D-F) were calculated with two-way ANOVA. ns: not significant. *: p<0.05; **: p<0.01; ***: p<0.001. Error bar: standard error of the mean. Each data point represents a single placenta or embryo.

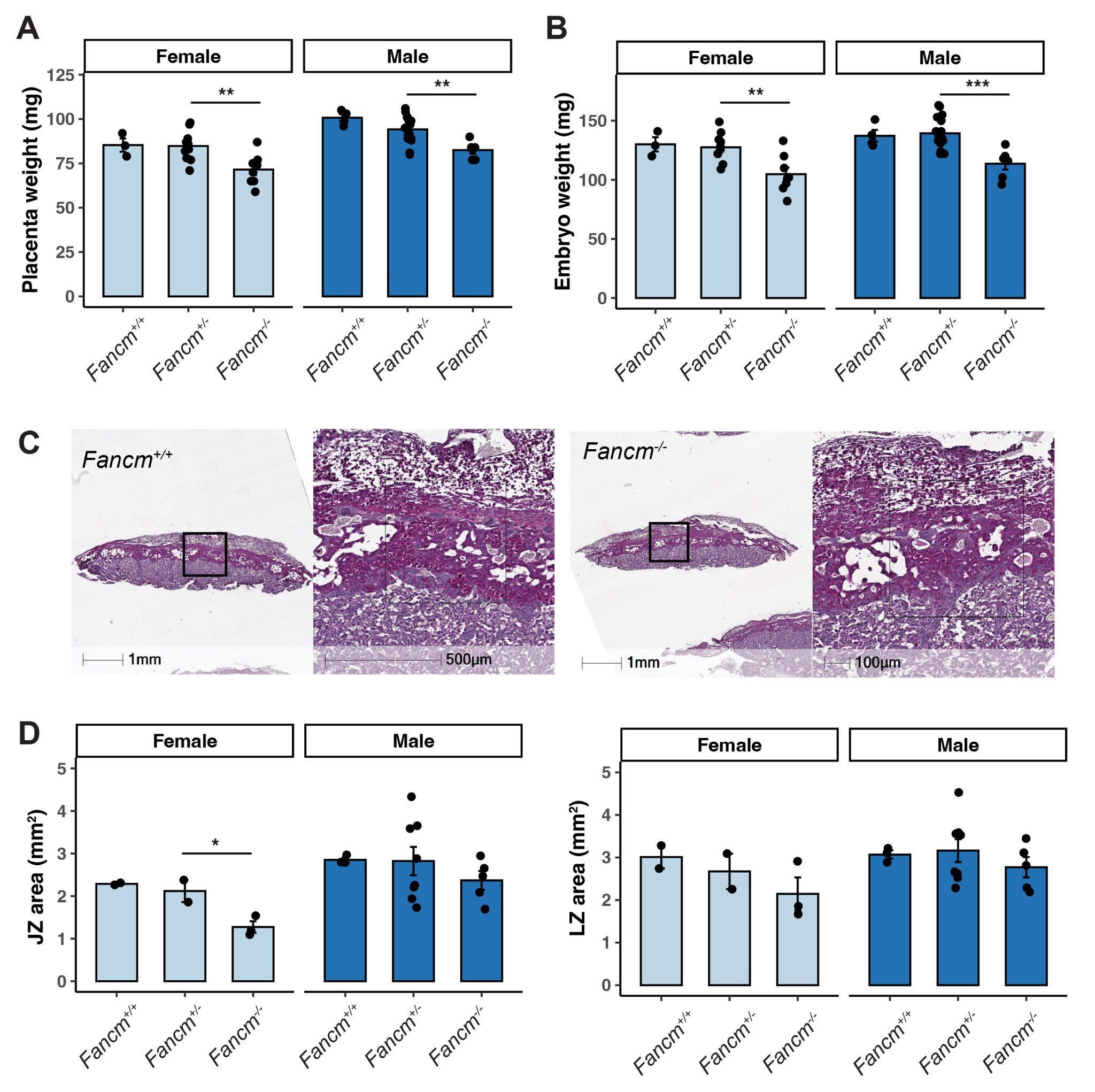

**Supplementary Figure 2. *Fancm* deficiency results in placental defect.** (A-B) Placental and embryonic weights of indicated genotypes at E13.5. (C) Periodic Acid Schiff staining of E13.5 placental sections from indicated genotypes. (D) Measurements of placental JZ and LZ areas. p-values were calculated with one-way ANOVA followed by Tukey’s HSD test. ns: not significant. *: p<0.05; **: p<0.01; ***: p<0.001. Error bar: standard error of the mean. Each data point represents a single placenta or embryo.

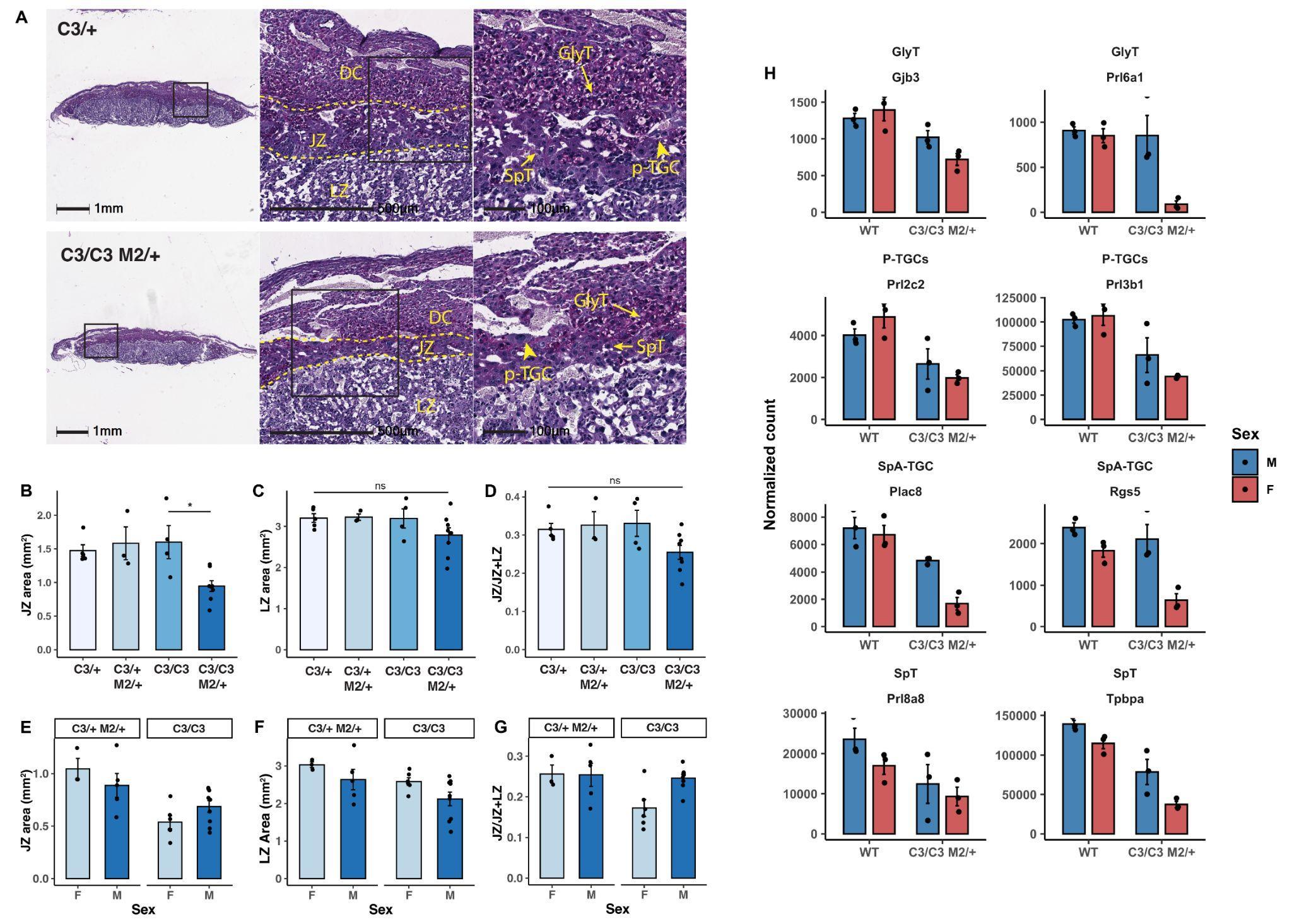

**Supplemental Figure 3. Placentae with replication stress have smaller JZ.** (A) Periodic Acid Schiff staining of placental sections from reciprocal matings at E13.5 for the indicated genotypes. (B-D) Measurements of placental JZ and LZ areas as well as the proportion of JZ from all indicated genotypes obtained from reciprocal matings at E13.5. (E-G) Comparison of JZ and LZ area as well as proportion of JZ in *Mcm4^C3/C3^ Mcm2^Gt/+^* genotype from sex-skewing and reciprocal matings. (H) Expression of trophoblast marker genes at E13.5 in WT and *Mcm4^C3/C3^ Mcm2^Gt/+^* placentae. WT: wild type; C3/+: *Mcm4^C3/+^*; C3/C3: *Mcm4^C3/C3^*; C3/+ M2/+: *Mcm4^C3/+^ Mcm2^Gt/+^*; C3/C3 M2/+: *Mcm4^C3/C3^ Mcm2^Gt/+^*. DC: decidua; JZ: junctional zone; LZ: labyrinth zone; p-TGC: parietal trophoblast giant cells; SpT: spongiotrophoblast; SynT: syncytiotrophoblast; GlyT: glycogen trophoblast; M: Male; F: Female. p-values were calculated with one-way ANOVA followed by Tukey’s HSD test. ns: not significant. *: p<0.05; **: p<0.01; ***: p<0.001. Error bar: standard error of the mean. Each data point in (B-G) represents the average measurement taken from at least three sections of the same placenta. Each data point in (H) represents the normalized count value from an individual placenta in the RNA-Seq experiment.

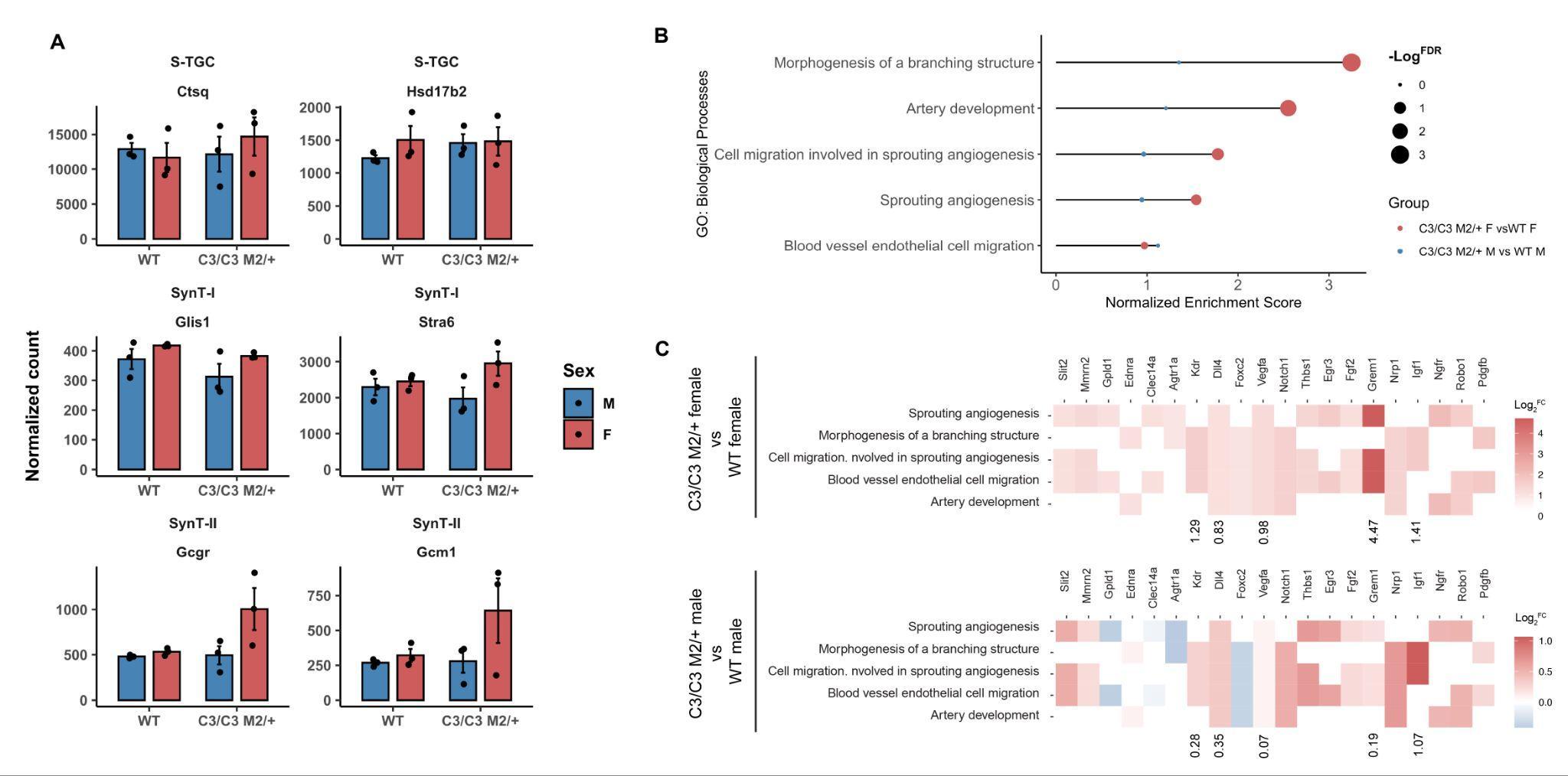

**Supplementary Figure 4. Placentae with replication stress have smaller junctional and labyrinth zones.** (A) Expression of trophoblast marker genes at E13.5 in WT and *Mcm4^C3/C3^ Mcm2^Gt/+^* placentae. (B) Gene enrichment analysis of GO terms related to angiogenesis. (C) Log transformed fold change of shared genes that are involved in angiogenesis in female and male mutant placentae compared to the WT respectively. WT: wild type; C3/C3 M2/+: *Mcm4^C3/C3^ Mcm2^Gt/+^*. S-TGC: sinusoidal trophoblast giant cells; SynT-I: syncytiotrophoblast I; SynT-III: syncytiotrophoblast II. Each data point in (A) represents the normalized count value from an individual placenta in the RNA-Seq experiment.

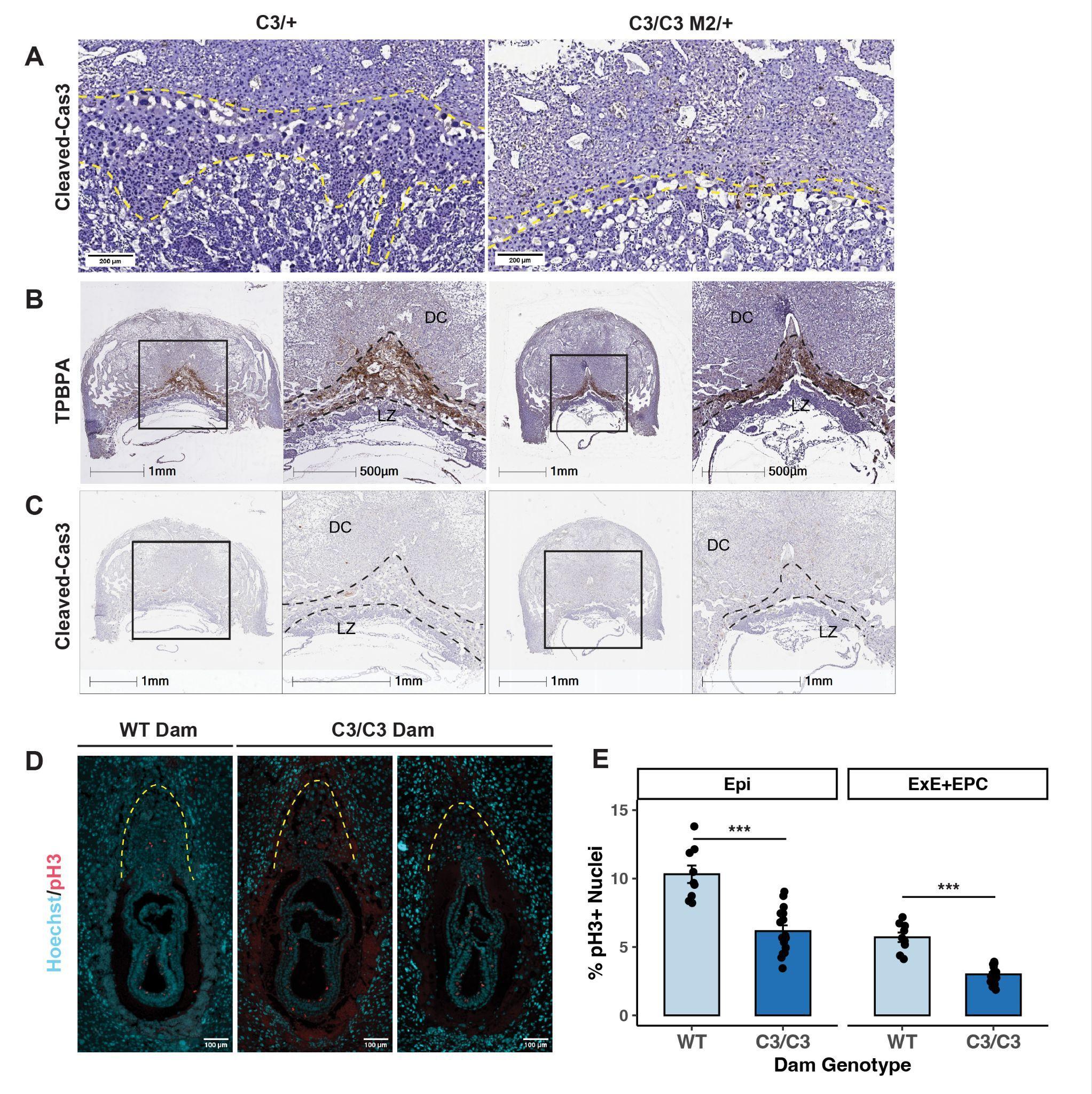

**Supplementary Figure 5. Placentae with replication stress have smaller junctional and labyrinth zones.** (A) Representative images of Cleaved-caspase 3 immunohistochemistry in E11.5 placental sections for the indicated genotypes. (B-C) Representative images of TPBPA and Cleaved-caspase 3 immunohistochemistry in E9.5 placental sections for the indicated genotypes. (D) Representative images of phospho-H3 immunofluorescence in E7.5 embryos sections. (E) Quantification of phospho-H3 positive cells in the epiblast and extraembryonic ectoderm. WT: wild type; C3/+: *Mcm4^C3/+^*; C3/C3: *Mcm4^C3/C3^*; C3/C3 M2/+: *Mcm4^C3/C3^ Mcm2^Gt/+^*. Epi: epiblast; ExE: extraembryonic ectoderm; EPC: ectoplacental cone; DC: decidua; LZ: labyrinth zone. Each data point in (E) represents the average percent positive cells from at least three consecutive sections of a single embryo.

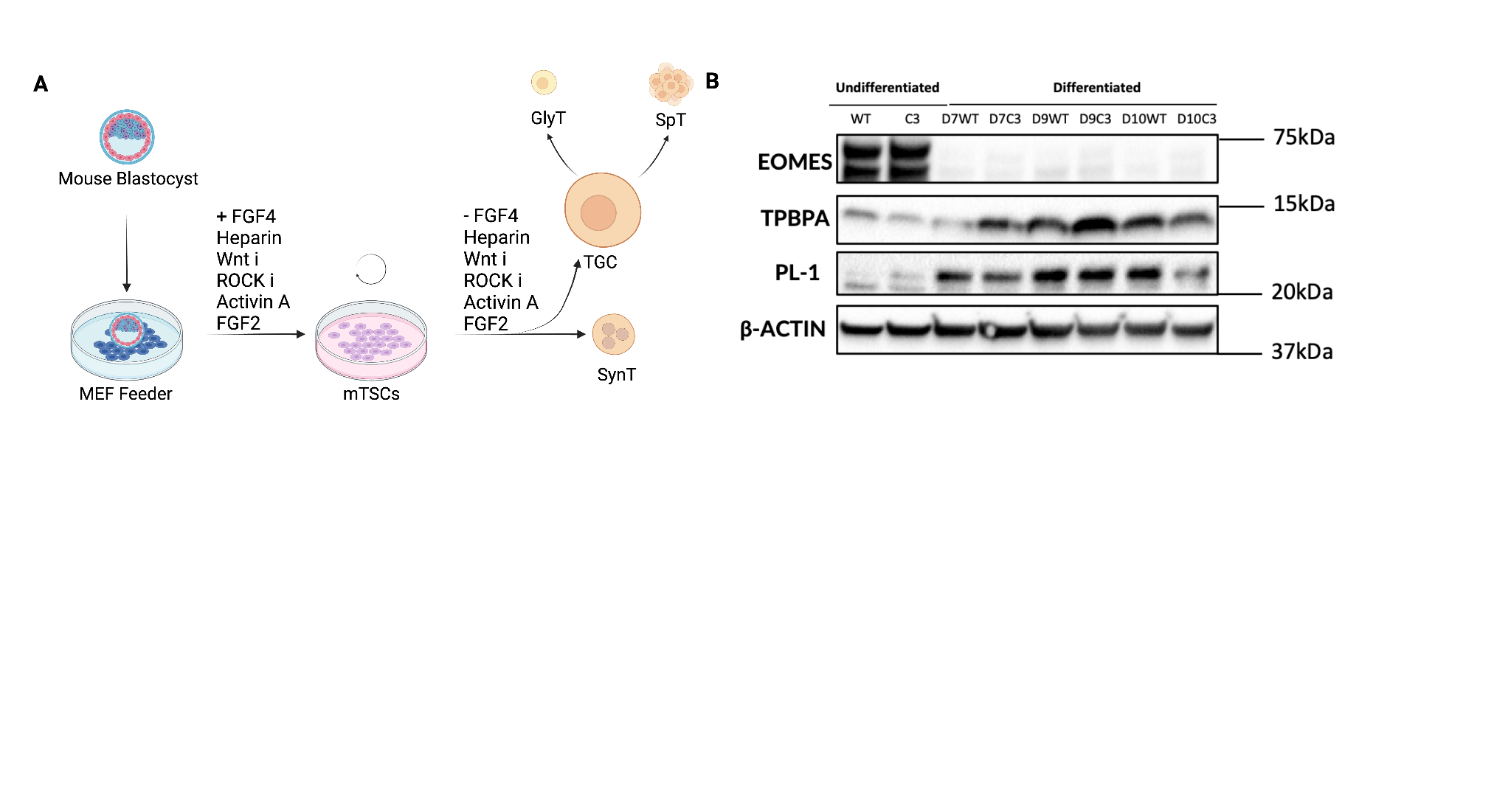

**Supplementary Figure 6: Trophoblast stem cell derivation procedure in defined medium and premature differentiation of *Chaos3* mutant TSCs.**

(A) Schematic representation of TSC derivation from E3.5 blastocysts under defined culture conditions. (B) Western blot analysis of trophoblast markers before and after TSC differentiation. Genotypes are indicated: C3, *Mcm4^Chaos3/Chaos3^*; D, day; PL-1 and TPBPA are markers of TGCs and spongiotrophoblast, respectively. β-actin is a loading control.

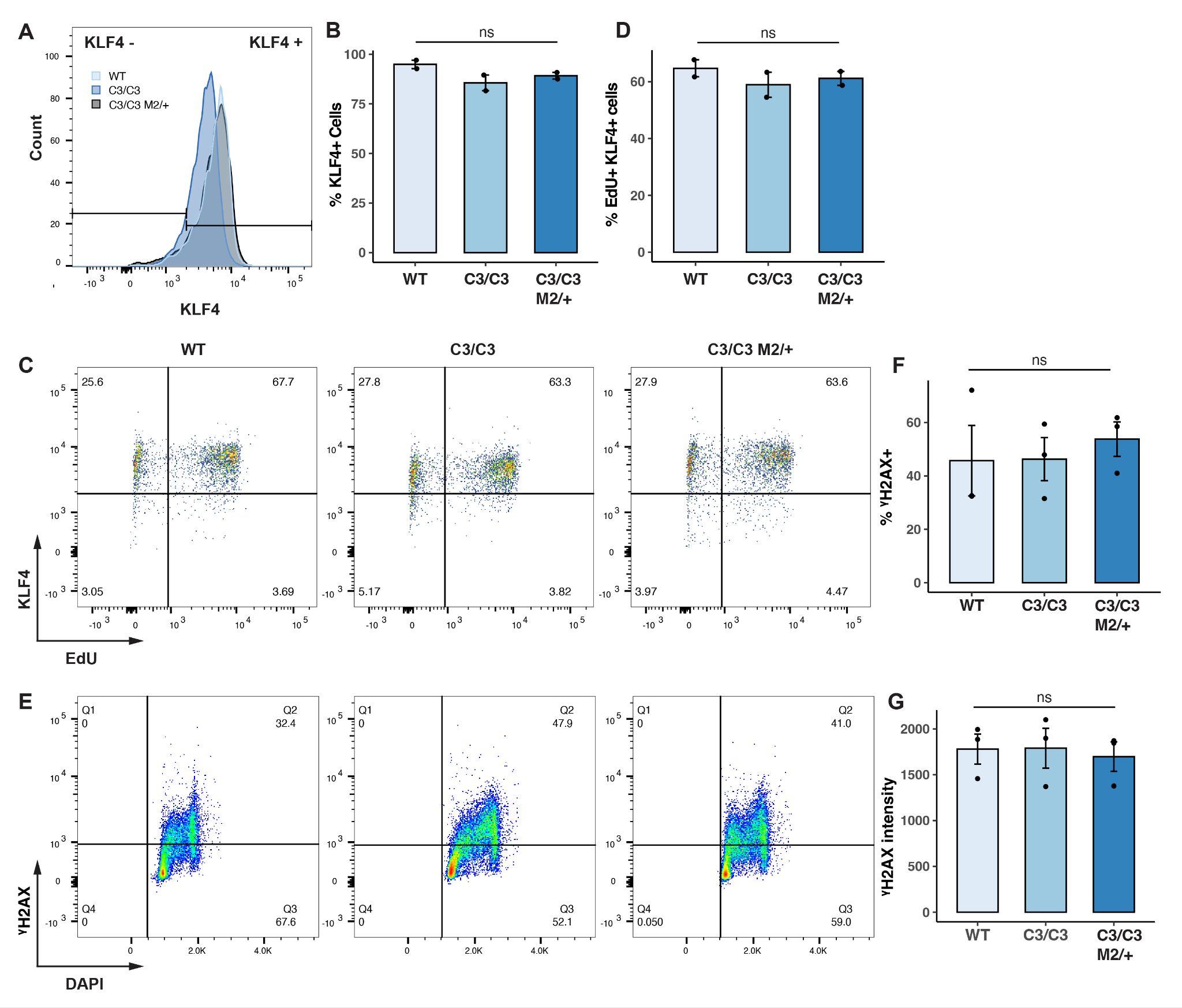

**Supplementary Figure 7: Mouse embryonic stem cells (ESCs) are not sensitive to RS induced by MCM2-7 deficiency.** (A) Flow cytometry analysis of naive pluripotency marker KLF4 in WT, C3/C3, and C3/C3 M2/+ ESCs. (B) Quantification of KLF4 positive cells in ESCs. (C) Flow cytometry analysis of EdU pulse labeled and KLF4 stained ESCs in indicated genotypes. (D) Quantification of the EdU and KLF4 double positive cells from flow cytometry. (E) Flow cytometry analysis of γH2AX in ESCs of the indicated genotypes. (F) and (G) Quantification of %positive γH2Ax cells and the average γH2Ax intensity in each indicated genotypes. Each data point in (B), (D), (F) and (G) represents biological replicates. Error bar: standard error of the means. ns: not significant; *p-values* were calculated using one-way anova.

**Supplementary Table 1**

| ♀*Mcm4^C3/C3^ Tmem173^gt/gt^ X* ♂*Mcm4^C3/+^ Mcm2^Gt/+^ Tmem173^gt/gt^* | | | | | |
| --- | --- | --- | --- | --- | --- |
|  | *Mcm4^C3/+^ Tmem173^gt/gt^* | *Mcm4^C3/+^ Mcm2^Gt/+^ Tmem173^gt/gt^* | *Mcm4^C3/C3^ Tmem173^gt/gt^* | *Mcm4^C3/C3^ Mcm2^Gt/+^ Tmem173^gt/gt^* | Total |
| Total | 42 | 51 | 23 | 2 | 118 |
| Male | 16 | 31 | 16 | 2 | 65 |
| Female | 26 | 20 | 7 | 0 | 53 |

*Chi-squared p-value = 2.0944E-10*

| ♀*Mcm4^C3/+^ Mcm2^Gt/+^ Tmem173^gt/gt^ X* ♂*Mcm4^C3/C3^ Tmem173^gt/gt^* | | | | | |
| --- | --- | --- | --- | --- | --- |
|  | *Mcm4^C3/+^ Tmem173^gt/gt^* | *Mcm4^C3/+^ Mcm2^Gt/+^ Tmem173^gt/gt^* | *Mcm4^C3/C3^ Tmem173^gt/gt^* | *Mcm4^C3/C3^ Mcm2^Gt/+^ Tmem173^gt/gt^* | Total |
| Total | 74 | 59 | 50 | 7 | 190 |
| Male | 38 | 35 | 24 | 4 | 101 |
| Female | 36 | 24 | 26 | 3 | 89 |

*Chi-squared p-value = 2.67332E-11*

| ♀*Mcm4^C3/C3^ Ddx58^-/-^ X* ♂*Mcm4^C3/+^ Mcm2^Gt/+^ Ddx58^-/-^* | | | | | |
| --- | --- | --- | --- | --- | --- |
|  | *Mcm4^C3/+^*  *Ddx58^-/-^* | *Mcm4^C3/+^ Mcm2^Gt/+^ Ddx58^-/-^* | *Mcm4^C3/C3^*  *Ddx58^-/-^* | *Mcm4^C3/C3^ Mcm2^Gt/+^ Ddx58^-/-^* | Total |
| Total | 49 | 40 | 41 | 10 | 140 |
| Male | 21 | 18 | 22 | 9 | 70 |
| Female | 28 | 22 | 21 | 1 | 72 |

*Chi-squared p-value = 1.40229E-05*

| ♀*Mcm4^C3/+^ Mcm2^Gt/+^ Ddx58^-/-^ X* ♂*Mcm4^C3/C3^ Ddx58^-/-^* | | | | | |
| --- | --- | --- | --- | --- | --- |
|  | *Mcm4^C3/+^ Ddx58^-/-^* | *Mcm4^C3/+^ Mcm2^Gt/+^ Ddx58^-/-^* | *Mcm4^C3/C3^ Ddx58^-/-^* | *Mcm4^C3/C3^ Mcm2^Gt/+^ Ddx58^-/-^* | Total |
| Total | 47 | 40 | 29 | 7 | 123 |
| Male | 29 | 21 | 14 | 7 | 71 |
| Female | 18 | 19 | 15 | 0 | 52 |

*Chi-squared p-value = 1.51087E-06*

**Supplementary Table 2**

| ♀*Mcm4^C3/+^ X* ♂*Mcm4^C3/+^* | | | | | |
| --- | --- | --- | --- | --- | --- |
| Condition | *Mcm4^+/+^* | *Mcm4^C3/+^* | *Mcm4^C3/C3^* | Total | *Chi-squared p-value* |
| Conventional | 36 | 22 | 5 | 63 | 1.3521E-08 |
| Defined | 18 | 35 | 5 | 58 | 0.015683087 |

**TSC derivation efficiency under conventional and defined culture conditions**. The number of successfully derived WT, *Mcm4^C3/+^* , and *Mcm4^C3/C3^* TSC lines is shown for two independent experiments. Each condition contains 2 independent experiments.

**Supplementary Table 3**

| **Gene** | **Placenta phenotype** (MGI annotations) |
| --- | --- |
| *Cdk7* | abnormal trophoblast layer morphology. |
| *Commd1* | abnormal extraembryonic tissue morphology; abnormal placenta vasculature; abnormal chorionic plate morphology. |
| *Cul4b* | abnormal extraembryonic tissue morphology; disorganized extraembryonic tissue; abnormal placenta labyrinth morphology; small placenta; pale placenta; disorganized placental labyrinth. |
| *Ddx11* | placental labyrinth hypoplasia; increased allantois apoptosis; failure of chorioallantoic fusion. |
| *Fzr1* | placental labyrinth hypoplasia; abnormal trophoblast giant cell morphology; pale placenta; abnormal placenta morphology; absent trophoblast giant cells; small placenta; abnormal placenta labyrinth morphology; abnormal placental labyrinth vasculature morphology; abnormal placenta size. |
| *Gtf2i* | failure of chorioallantoic fusion. |
| *Hus1* | abnormal placenta labyrinth morphology; abnormal placental labyrinth vasculature morphology; failure of chorioallantoic fusion. |
| *Kat7* | failure of chorioallantoic fusion. |
| *Mapk14* | abnormal placental labyrinth vasculature morphology; abnormal placenta morphology; decreased trophoblast giant cell number; absent placental labyrinth; decreased spongiotrophoblast size; abnormal placenta development; abnormal placenta labyrinth morphology; abnormal trophoblast layer morphology; abnormal placenta vasculature; impaired placental function; decreased placental labyrinth size. |
| *Myc* | abnormal placenta morphology; placental labyrinth hypoplasia; abnormal trophoblast layer morphology. |
| *Ncoa6* | abnormal placenta morphology; abnormal placenta vasculature; abnormal placental labyrinth vasculature morphology; decreased spongiotrophoblast size; abnormal placenta development; abnormal trophoblast giant cell morphology; absent placental labyrinth. |
| *Nipbl* | abnormal placenta labyrinth morphology; increased placenta weight; abnormal placenta junctional zone morphology; abnormal spongiotrophoblast cell morphology; abnormal placenta physiology. |
| *Npm1* | small placenta; abnormal placenta development. |
| *Pagr1a* | abnormal extraembryonic tissue morphology; increased allantois apoptosis; abnormal extraembryonic ectoderm morphology. |
| *Rad9a* | abnormal chorioallantoic fusion. |
| *Rtel1* | decreased trophoblast giant cell number; small ectoplacental cone; abnormal placenta vasculature; failure of chorioallantoic fusion; abnormal placenta morphology. |
| *Senp2* | absent trophoblast giant cells; abnormal trophoblast layer morphology; abnormal trophoblast giant cell morphology; small placenta; decreased spongiotrophoblast size. |
| *Setd2* | abnormal placenta labyrinth morphology; abnormal placental labyrinth vasculature morphology; failure of chorioallantoic fusion. |
| *Smchd1* | abnormal trophoblast layer morphology; abnormal trophoblast giant cell morphology. |
| *Stk11* | abnormal spongiotrophoblast layer morphology; small placenta; abnormal placental labyrinth vasculature morphology; delayed chorioallantoic fusion; placenta hemorrhage; abnormal placenta development. |
| *Trip12* | abnormal placenta development; abnormal placenta labyrinth morphology. |
| *Ubr5* | failure of chorioallantoic fusion. |
| *Wrap53* | abnormal placenta morphology; abnormal trophoblast layer morphology. |
| *Ehmt2* | decreased trophoblast giant cell number; small placenta. |
| *Jun* | abnormal placenta morphology. |
| *Zpr1* | abnormal trophoblast giant cell morphology. |

**Supplementary Table 4 – Guide and Primer sequences**

| **Primer name** | **Gene** | **Sequence (5’ to 3’)** |
| --- | --- | --- |
| Chaos3_gF2 | Mcm4 | CATTAACAAAACCAACAGATA |
| Chaos3_gR2 | Mcm4 | TTAGCGGCATACCCAGGC |
| Mcm2_gF4 | Mcm2 | TCCACTCTGATGGGCTGT |
| Mcm2_GTgR4 | Mcm2 | ATGTATGCTATACGAACGGTAGGAT |
| Sry_R | Sry | CCACTCCGTGACACTTTAGCCCTCCGA |
| Sry_L | Sry | TTGTCTAGAGAGCATGGAGGGCCATGTCAA |
| RR0002-R | Mcm3 | GGATGAGGGAGCAGGGCTCGGCAC |
| RR0002-F | Mcm3 | CACTGTTTATATGTGCACGTGTACC |
| RR0002-WR | Mcm3 | CTTCTGTCGCTTTCAGACCAGAAGC |
| Myd88_mut_gF1 | Myd88 | CCACCCTTGATGACCCCCTA |
| Myd88_gF1 | Myd88 | GTTGTGTGTGTCCGACCGT |
| Myd88_gR1 | Myd88 | GTCAGAAACAACCACCACCATGC |
| Tmem173_gF2 | Tmem173 | GGGGCAGCATATCTCGGAAT |
| Tmem173_gR2 | Tmem173 | AGAAGGCTAACGAGCTGAGTG |
| Ddx58_gF1 | Ddx58 | TGGACTTTGTGAAGCCATCGAA |
| Ddx58_gR1 | Ddx58 | ACCTGGCATTGGTTTACCTGTC |

**Supplementary Table 5 – Antibody and cell labeling reagents**

| **Primary Antibodies** | **Dilutions** | **Supplier** | **Catalog #** |
| --- | --- | --- | --- |
| Rb Anti-Ki67 | 1:250 | Cell Signaling Technology | 12202S |
| Rb Anti-Cleaved Caspase 3 | 1:250 | Cell Signaling Technology | 9661S |
| Rb Anti-EOMES | 1:500 | Abcam | ab23345 |
| Ms Anti-CDX2 | 1:500 | BioGenX | AM392-5M |
| Rb Anti-PL1 | 1:250 | Abcam | ab15554 |
| Rb Anti-TPBPA | 1:500 | Abcam | ab104401 |
| Rb Anti-MCT4 | 1:250 | Sigma-Aldrich | AB3314P |
| Ch Anti-MCT1 | 1:250 | Sigma-Aldrich | AB1286-I |
| Ms Anti-phospho H3 (Ser10) | 1:250 | Cell Signaling Technology | 9706S |
| Ms Anti-beta-ACTIN | 1:5000 | Sigma-Aldrich | A1978 |
| Ms Anti- CDK1 | 1:500 | Thermo Fisher | MA5-11472 |
| Rb Anti-Phospho-CDK1(Thr 14/15) | 1:1000 | Thermo Fisher | 44-686G |
| Rb Anti-Geminin | 1:1000 | Santa Cruz Biotechnology | SC-13015 |
| Rb Anti- CHK1 | 1:1000 | Santa Cruz Biotechnology | SC-7898 |
| Rb Anti-Phospho-CHK1 (Ser345) | 1:1000 | Cell Signaling Technology | 2348 |
| Rb Anti-Phospho-P53 (Ser15) | 1:1000 | Cell Signaling Technology | 12571 |
| Rb Anti-P53 | 1:1000 | Cell Signaling Technology | 32532 |
| Rb Anti-𝛄H2Ax (Ser139) | 1:1000 | Cell Signaling Technology | 9718 |
| Rb Anti-P21 (Waf1/Cip1) | 1:1000 | Cell Signaling Technology | 64016 |
| Ms Anti-KLF4(1E5) | 1:250 | Invitrogen | MA5-15672 |
| Secondary Antibodies | Dilutions | Supplier | Catalog # |
| Goat Anti-Rabbit Alexa Fluor 594 | 1:500 | Thermo Fisher | A-11012 |
| Goat Anti-Chicken Alexa Fluor 488 | 1:500 | Thermo Fisher | A-11039 |
| Goat Anti-Mouse Alexa Fluor 594 | 1:500 | Thermo Fisher | A-11032 |
| Goat Anti-Mouse IgG (H+L), HRP | 1:5000 | Thermo Fisher | 31430 |
| Goat Anti-Rabbit IgG (H+L), HRP | 1:2000 | Thermo Fisher | 31460 |
| DAPI | 1:5000 | Thermo Fisher | D1306 |
